# Analysis and annotation of genome-wide DNA methylation patterns in two nonhuman primate species using the Infinium Human Methylation 450K and EPIC BeadChips

**DOI:** 10.1101/2020.05.09.081547

**Authors:** Fabien Pichon, Florence Busato, Simon Jochems, Beatrice Jacquelin, Roger Le Grand, Jean-Francois Deleuze, Michaela Müller-Trutwin, Jörg Tost

**Affiliations:** Laboratory for Epigenetics and Environment, Centre National de Recherche en Génomique Humaine, CEA-Institut de Biologie François Jacob, Evry, France; Institut Pasteur, HIV Inflammation and Persistence Unit, Paris, France; Université Paris Diderot, Sorbonne Paris Cité, Paris, France; CEA, Université Paris Sud, INSERM U1184, Immunology of Viral Infections and Autoimmune Diseases (IMVA), IDMIT Department / IBFJ, Fontenay-aux-Roses, France

**Author notes:** **Corresponding author** Jörg Tost, Laboratory for Epigenetics and Environment, Centre National de Recherche en Génomique Humaine, CEA-Institut de Biologie François Jacob, Evry, France.

**Keywords:** Infinium, 450K, EPIC, microarray, DNA methylation, macaca, chlorocebus, rhesus macaque, monkey, African green monkey, vervet, annotation

## Abstract

The Infinium Human Methylation450 and Methylation EPIC BeadChips are useful tools for the study of the methylation state of hundreds of thousands of CpG across the human genome at affordable cost. However, in a wide range of experimental settings in particular for studies in infectious or brain-related diseases, human samples cannot be easily obtained. Hence, due to their close developmental, immunological and neurological proximity with humans, non-human primates are used in many research fields of human diseases and for preclinical research. Few studies have used DNA methylation microarrays in simian models. Microarrays designed for the analysis of DNA methylation patterns in the human genome could be useful given the genomic proximity between human and nonhuman primates. However, there is currently information lacking about the specificity and usability of each probe for many nonhuman primate species, including rhesus macaques (*Macaca mulatta*), originating from Asia, and African green monkeys originating from West-Africa (*Chlorocebus sabaeus)*. Rhesus macaques and African green monkeys are among the major nonhuman primate models utilized in biomedical research. Here, we provide a precise evaluation and re-annotation of the probes of the two microarrays for the analysis of genome-wide DNA methylation patterns in these two Cercopithecidae species. We demonstrate that up to 162,000 of the 450K and 255,000 probes of the EPIC BeadChip can be reliably used in *Macaca mulatta* or *Chlorocebus sabaeus*. The annotation files are provided in a format compatible with a variety of preprocessing, normalization and analytical pipelines designed for data analysis from 450K/EPIC arrays, facilitating high-throughput DNA methylation analyses in *Macaca mulatta* and *Chlorocebus sabaeus*. They provide the opportunity to the research community to focus their analysis only on those probes identified as reliable. The described analytical workflow leaves the choice to the user to balance coverage versus specificity and can also be applied to other Cercopithecidae species.

## Background

DNA methylation is an epigenetic mark associated with gene regulation. It impacts a number of key biological processes including genomic imprinting, X-chromosome inactivation, repression of transposable elements, aging, carcinogenesis and immunity against infectious diseases [1]. DNA methylation consists in the methylation of a cytosine (5-methylcytosine) mostly immediately followed by a guanine. Because of this methylation, cytosines are susceptible to deamination yielding thymines [2–4]. Due to their increased mutation rate, CpG dinucleotides have been depleted during evolution and are thus under-represented in the genome [5]. However, a higher density of mostly unmethylated CpG dinucleotides is found in CpG islands, generally localized in the first exon and intron or in the promoter region of genes [6,7].

CpG methylation can be measured in humans using the Infinium Human Methylation450 BeadChip Array (Infinium 450K), which measures methylation levels at more than 450,000 CpG across the human genome [8]. This array has been replaced by the Infinium Human MethylationEPIC BeadChip Array (Infinium EPIC, which adds about 350,000 CpGs localized in enhancers; [9]). Due to its accuracy and capacity to analyze large cohorts at an affordable cost, the Infinium 450K microarray has been used in a wide range of epigenome-wide association studies in humans (see review in [10]).

Only few genome-wide array tools are currently available for nonhuman primate models. Of note, these species have a close phylogenic proximity with humans and a high percentage of DNA identity and gene homology, raising the possibility of the use of human microarrays in studies in non-human primate models. Indeed, human gene expression microarrays have been used in various studies on monkeys, from HCV and SIV infection [11–13] to asthma [14] or glaucoma [15]. More recently, the human Affymetrix HG-U133 Plus 2.0 GeneChip has also been used for a gene expression study in the phylogenetically more distant lemur *Microcebus murinus*, which has been shown to be an excellent model for Alzheimer’s disease ([16]; Pichon et al., unpublished data). In the latter study, human microarrays detected about 20% of lemur transcripts, which is expected because of the divergence of both species.

Similarly, the human Infinium 450K was also used for DNA methylation studies in great apes [17]. Seventy-three percent of the probes designed to the human genome mapped to the bonobo genome, 72%-77% to chimpanzee, 61% to gorilla and 44% to orangutan genomes [17,18]. Studies were also performed in monkeys of the cynomolgus macaque species (*Macaca fascicularis*) [19], rhesus macaque (*Macaca mulata, MM*) [18] and baboon [20]. For example, using different selection criteria for probes yielding reliable signals, 61% of human probes were mapped and annotated to the *Macaca fascicularis* genome and subsequently used to study the impact of birth weight on gene methylation and expression in *Macaca fascicularis* [21]. Microarrays initially designed for the interrogation of the human genome have thus been used to study gene expression or DNA methylation of nonhuman primate samples. This can be successfully done under the condition that a thorough evaluation of reliable CpG targeting probes is performed for the species of interest.

Several monkey species are widely used models to study complex human diseases contributing to unraveling the mechanisms or treatment of human and animal diseases. Rhesus macaques are for instance frequently used for the development of vaccines against viral diseases, including Sars-CoV-2, HIV, Influenza and Ebola virus [22–24]. They are also widely used in various studies ranging from development and imprinting, to addiction and social cognition [25–30]. In parallel, the *Chlorocebus* genus, among which figure prominently African green monkeys (AGM), have been a gold standard model for the studies and fight against several infectious diseases, such as yellow fever, trypanosoma and plague in the past [31,32]. African green monkeys, in particular *Chlorocebus sabaeus*, are included in studies of neurological disorders, in pharmacological trials [33] and more recently also for the identification of mechanisms of protection against HIV/AIDS, MERS-CoV and SARS-CoV2 [34–37].

*Macacca mulatta* and *Chlorocebus* have therefore, together with baboons and cynomolgus macaques, become reference animal models for preclinical research. Nonetheless, no commercial off-the-shelf DNA methylation microarrays are available for these species. In the present study, we evaluated the use of the Infinium 450K and Infinium EPIC BeadChips for genome-wide DNA methylation analyses in samples from *Chlorocebus sabaeus* and *Macaca mulatta* and conducted an in-depth analysis of the available probes for these two old world monkey genomes. Results show that about one third of the Infinium 450K or EPIC human-designed probes can be reliably used to study DNA methylation in these Cercopithecidae and that the majority map to gene features. We provide for each species and each microarray a list of annotated probes that can be used by the scientific community for genome-wide DNA methylation studies in *Chlorocebus sabaeus* or *Macaca mulatta.* These detailed data on probe behavior were obtained with stringent criteria. They provide the flexibility to the research community to focus their analysis only to those probes identified as reliable.

## Methods

### Study approval

Animals were housed at the IDMIT center of the Commissariat à l’Energie Atomique (CEA, Fontenay-aux-Roses, France, permit number: A 92-032-02). The CEA complies with Standards for Human Care and Use of Laboratory of the Office for Laboratory Animal Welfare (OLAW, USA) under OLAW Assurance number #A5826-01. The Central Committee for Animals at Institut Pasteur or Ethical Committee of Animal Experimentation (CETEA-DSV, IDF, France) approved all animal experimental procedures (Notification numbers: 10-051b, 12-006b). The studies were conducted in strict accordance with the international European guidelines 2010/63/UE on protection of animals used for experimentation and other scientific purposes (French decree 2013-118).

### Samples

Blood was collected from 13 AGM (*Chlorocebus sabaeus*) and 17 rhesus macaques (*Macaca mulatta*) by venipuncture on EDTA tubes. As several blood samples were available for several animals, in total 21 *Chlorocebus sabaeus* and 25 *Macaca mulatta* samples were included in the study. CD4+ peripheral blood mononuclear cells (PBMC) were purified, as described previously using magnetic anti-CD4 beads (Miltenyi)[13]. CD4+ T cell purity after isolation was confirmed using flow cytometry (median 97%, IQR 93%-98%). DNA was extracted from CD4+ T cells using the DNeasy blood and tissue kit (Qiagen), according to manufacturer’s protocol.

### Infinium 450K analyses

One µg of DNA was bisulfite-treated using the EpiTect® 96 Bisulfite Kit (Qiagen, Hilden, Germany) and analysed using the Infinium Human Methylation 450K BeadChips (Illumina, San Diego, CA) according to the manufacturer’s protocol.

### Mapping of 50 bp probe sequences from Infinium 450K and EPIC

References genomes used in this work were downloaded from Ensembl: MMUL1.0 (corresponding to UCSC rheMac2) for *Macaca mulatta* and ChlSab1.1 for *Chlorocebus sabaeus*. Infinium 450K microarrays provide DNA methylation measures for 482,421 CpG sites across the human genome (135,476 Infinium I and 346,945 Infinium II probes). In the same way, Infinium EPIC allows measuring of 863,904 CpG (142,262 Infinium I and 721,642 Infinium II probes. 14,870 (3.1%) probes of the Infinium 450K and 18,903 (2.2%) probes of the Infinium EPIC contain two CpG separated by exactly 46 base pairs (bp), and thus located at each probe sequence extremity. In such cases, we used Blat [38] to determine which CpG was actually measured by the probe.

We then mapped all the 50 bp probe sequences targeting CpG positions in the human genome from the Infinium 450K and EPIC arrays to *Chlorocebus sabaeus (CS)* and *Macaca mulatta* (*MM*) genomes, consecutively, using Bowtie [39], allowing only a unique position on the respective genome and up to 3 mismatches. Sequences were thus classified as “Perfect Match”, “1 mismatch”, “2 mismatches”, “3 mismatches” or “Unmapped” depending on the number of mismatches attributed by Bowtie. Unmapped probes included thus probes with either more than 3 mismatches as well as those that could not be mapped to unique location. Because of the necessity to only keep probe sequences that can reliably hybridize on simian genomes and inform on methylation state of the CpG sites, we removed probes containing mismatches at the CpG site, which were annotated as “CS-non-targeting” or “MM-non-targeting”. Sequences with intact CpG sites were qualified as “CS-functional” or “MM-functional”, for *Chlorocebus sabaeus* and *Macaca mulatta* respectively. Among the CS-functional or MM-functional probes, we also determined the exact position of the closest mismatch, if any, relative to the CpG if the mismatch was localized from 1 bp to 10 bp away from the CpG to yield the final selction of valid probes.

### Annotation of Infinium 450K and EPIC probes in *Macaca mulatta* and *Chlorocebus sabaeus*

To annotate probes, the annotations files for *Macaca mulatta* and *Chlorocebus sabaeus* were retrieved from Ensembl (Version 84), containing respectively 44,725 and 28,078 transcripts for 30,246 and 27,985 genes (coding, non-coding and pseudogenes). In accordance with the annotation file provided by Illumina for the arrays we divided transcripts into 6 different categories: promoter region ranging from 1 bp to 200 bp upstream of the TSS (TSS200), promoter region ranging from 201 bp to 1,500 bp upstream of the TSS (TSS1500), 5’ untranslated region (5’UTR), first exons (1stExon), 3’ untranslated region (3’UTR) and gene bodies, excluding the 5’ and 3’ UTRs and first exons (Body). Similarly, CpG islands prediction files were downloaded from the UCSC genome browser (containing respectively 24,226 and 26,663 CpG islands for *Macaca mulatta* and *Chlorocebus sabaeus*) for the annotation relative to CpG islands following again the annotation criteria used by Illumina for the human genome. UCSC islands predictions are based on the following parameters: CpG obs/exp ratio > 0.6, CG content > 50%, length > 200bp. Shores are defined as island flanking regions ranging to up to 2,000 bp and shelves are defined as island flanking regions ranging from 2,001 bp to 4,000 bp. Northern and southern shores and shelves (noted N_ and S_) are respectively defined as the upstream and downstream shores or shelves according to chromosomal coordinates.

### Preprocessing and correction for Infinium I/II shift of 450K data for monkeys

To normalize and identify differentially methylated probes in the two monkey species, a refined version of subset quantile normalization (SQN) pipeline [40], which performs the SQN at the level of each individual sample prior to a between sample quantile normalization, was used to correct for the difference in the performance and dynamic range of Infinium I and Infinium II probes. The original Illumina annotation file used in the pipeline were replaced by the ones created for *Macaca mulatta* and *Chlorocebus sabaeus*. Due to the stringent criteria in the probe selection process, no further filtering for e.g. non-specific probes was required.

### Pyrosequencing analysis

Quantitative DNA methylation analysis for validation was performed by pyrosequencing of bisulfite-treated DNA [41]. Six regions of interest for validation were amplified using 30 ng of bisulfite-treated human genomic DNA and 5 to 7.5 pmol of forward and reverse primer, one of them being biotinylated. Sequences for oligonucleotides for PCR amplification and pyrosequencing are shown in Supplementary Table 1. Reaction conditions were either 1x HotStar Taq buffer supplemented with 1.6 mM MgCl_2_, 100 µM dNTPs, 5 pM of each primer and 2.0 U HotStar Taq polymerase (Qiagen) in a 25 µl volume or 1X Phusion U Hotstart (Thermo Fisher Scientific) with 5 pM of each primer. The PCR program consisted of a denaturing step of 15 min at 95°C followed by 50 cycles of 30 s at 95°C, 30 s at the respective annealing temperature and 20 s at 72°C, with a final extension of 5 min at 72°C. 10 µl of PCR product were rendered single-stranded as previously described [41] and 4 pmol of the respective sequencing primer were used for analysis. Quantitative DNA methylation analysis was carried out on a PSQ 96MD system with the PyroGold SQA Reagent Kit (Qiagen) and results were analyzed using the PyroMark CpG software (V.1.0.11.14, Qiagen).

## Results

Throughout the manuscript, we use a nomenclature of “targeting” probes, referring to probes that map and potentially target a CpG dinucleotide in the respective simian genomes allowing still for mismatches at any place in the probe including the CpG site, and “functional probes”, which target an intact CpG site in the simian genomes, as well as “valid probes”, which is the final selection of probes based on the selection criteria described below in the results section. First, we mapped the human probes to the simian genomes. All 50 bp probe sequences designed to target a CpG in the human genome, were extracted from Illumina’s manifest files for each microarray (482,421 sequences for the Infinium 450K and 863,904 for the Infinium EPIC).

### Mapping probes of the Infinium 450K and EPIC BeadChip on the Chlorocebus sabaeus genome

From the 482,421 Infinium 450K CpG probes designed for the human genome, 231,217 (47.9%) mapped to *Chlorocebus sabaeus* (CS) genome according to our parameters, of which 36,439 (15.8%) were perfect matches, 57,785 (25.0%) had one mismatch, 69,312 (30.0%) had two and 67,681 (29.3%) had three mismatches (Figure 1, Table 1). A more careful study of the mismatch position showed that 59,305 (25.6%) probe sequences had at least one mismatch at the CpG site and were thus identified as CS-non-functional probes. Of note, these non-targeting probes presented a total of 60,377 mismatches of which 71.3% were T➔C or A➔G, *i.e.* weak to strong (W➔S), nucleotide substitutions (*see* Supplementary Table 2). Out of the 231,217 mapped probes, 171,912 (74.4%) probe sequences were targeting a CpG dinucleotide and were thus identified as CS-functional (Table 1). From these CS-functional probes, 21.2.% has no mismatch at all, 51.7 % had the mismatch at more than 10 bp from the CpG and 46,457 (27 %) presented at least one mismatch at 10 bp or less from the CpG (Supplementary Table 3). The latter were further analyzed in more detail as described below.

**Figure 1:**
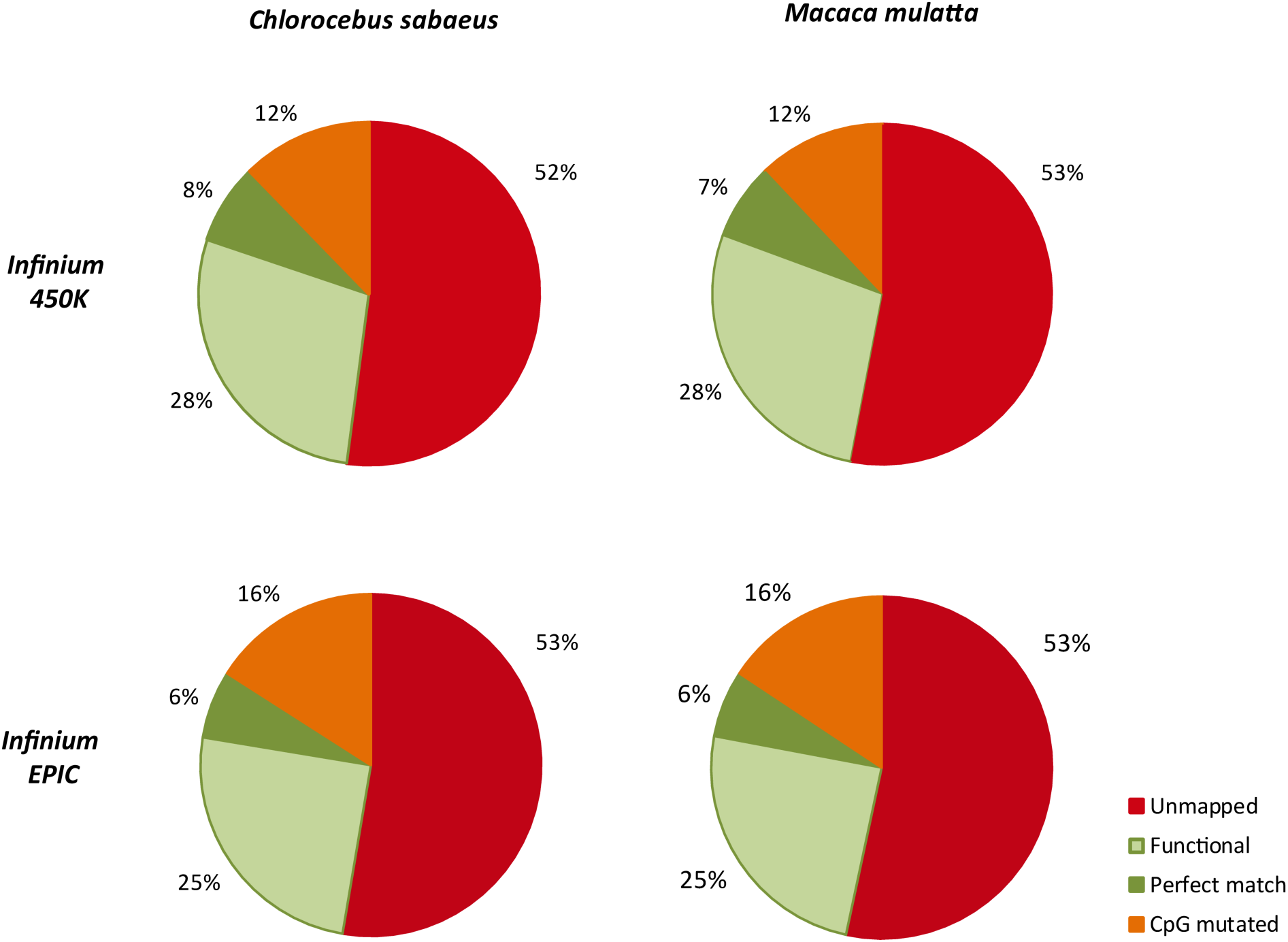
Proportion of probes of the Infinium 450K and the EPIC BeadChip mapping perfectly, with mismatches (functional), with a mutation at the CpG nucleotide or with more than three mismatches (unmapped) to the *Chlorocebus sabaeus* and *Macaca mulatta* genome.

**Table 1:** Mapping of the 50 bp probe sequences from the Infinium 450K and EPIC microarrays on the two simian genomes.

| Methylation array | Infinium 450K |  |  |  | Infinium EPIC |  |  |  |
| --- | --- | --- | --- | --- | --- | --- | --- | --- |
|  | *Chlorocebus sabaeus* |  | *Macaca mulatta* |  | *Chlorocebus sabaeus* |  | *Macaca mulatta* |  |
| Reference genome |  |  |  |  |  |  |  |  |
| Total CpG probes | 482,421 | 100.0% | 482,421 | 100.0% | 863,904 | 100.0% | 863,904 | 100.0% |
| Unmapped | 251,204 | 52.1% | 255,766 | 53.0% | 454,971 | 52.7% | 460,969 | 53.4% |
| Targeting | 231,217 | 47.9% | 226,655 | 47.0% | 408,933 | 47.3% | 402,935 | 46.6% |
| 0 mismatch in targeting | 36,439 | 15.8% | 35,386 | 15.6% | 55,479 | 13.6% | 54,789 | 13.6% |
| 1 mismatch in targeting | 57,785 | 25.0% | 57,257 | 25.3% | 97,668 | 23.9% | 97,505 | 24.2% |
| 2 mismatches in targeting | 69,312 | 30.0% | 68,226 | 30.1% | 126,889 | 31.0% | 125,108 | 31.0% |
| 3 mismatches in targeting | 67,681 | 29.3% | 65,786 | 29.0% | 128,897 | 31.5% | 125,533 | 31.2% |
| Functional in targeting | 171,912 | 74.4% | 168,466 | 74.3% | 270,922 | 66.3% | 267,237 | 66.3% |
| Functional 0 mismatch | 36,439 | 21.20% | 35,386 | 21.00% | 55,479 | 20.48% | 54,789 | 20.50% |
| Functional 1 mismatch | 47,691 | 27.74% | 47,350 | 28.11% | 75,028 | 27.69% | 74,938 | 28.04% |
| Functional 2 mismatches | 47,831 | 27.82% | 46,913 | 27.85% | 76,717 | 28.32% | 75,361 | 28.20% |
| Functional 3 mismatches | 39,951 | 23.24% | 38,817 | 23.04% | 63,698 | 23.51% | 62,149 | 23.26% |
| Non-functional in targeting | 59,305 | 25.6% | 58,189 | 25.7% | 138,011 | 33.7% | 135,698 | 33.7% |

We detailed the distribution of all the substitutions from the entire mapped probes set (Supplementary Table 4). We observed that a nucleotide located in a CpG site had more chances (around 13%) to be substituted than a base located outside a CpG site (about 3%). Moreover, about 60% of these substitutions are W➔S substitutions in CpG sites, while they represented around 20% along the whole 50 bp probe sequence.

Results from the Infinium EPIC BeadChip were very similar to those from Infinium 450K, with 408,933 (47.3%) CpG probes designed for the human genome mapping to the *Chlorocebus sabaeus* genome. Of these, 55,479 (13.6%) were perfect matches, 97,668 (23.9%) had one mismatch, 126,889 (31.0%) had two mismatches and 128,897 (31.5%) had three mismatches. From these mapped probes, 270,922 (66.3%, *versus* 74,4% in Infinium 450K) were identified as CS-functional and 138,011 were identified as CS-non-functional (Table 1). From these CS-functional probes, 73,853 presented at least one mismatch at 10 bp or less from the CpG (Supplementary Table 3) and were further investigated as described below.

As for Infinium 450K, we analyzed the distribution of all the substitutions from the whole mapped probe set (Supplementary Tables 5). From this distribution, it appeared that a base located in a CpG sites had higher chance to be substituted on the EPIC BeadChip (around 17%) than for Infinium 450K (around 13%, hypergeometric test p<0.001). In CpG sites, W➔S nucleotide substitutions represented around 65%, while they represented around 20% along the entire 50 bp probe sequences.

Altogether, for their use in DNA methylation analysis using the two human BeadChips, 31.4 (EPIC) and 35.6 % (450K) of the probes could be classified as functional for *Chlorocebus sabaeus*.

### Mapping probes of the Infinium 450K and EPIC BeadChip on the Macaca mulatta genome

Similarly to *Chlorocebus saba*eus, we found that of the 482,421 Infinium 450K CpG probes designed for the human genome, 226,655 (47.0%) mapped to the *Macaca mulatta* genome. Of these, 35,386 (15.6%) were perfect matches, 57,257 (25.3%) had one mismatch, 68,226 (30.1%) had two and 65,786 (29.0%) had three mismatches. Investigating the position of the respective mismatches showed that 58,189 (25.7%) probe sequences designed to the human genome had at least one mismatch at the CpG site and were thus identified as MM-non-functional (representing 59,228 mismatches) among which 71.2% were W➔S substitutions (Supplementary Figure 1, Supplementary Table 3). 168,466 (74.3%) probe sequences designed to the human genome were identified as MM-functional (Table 1). From these MM-functional probes, 45,462 presented at least one mismatch at 10 bp or less from the CpG (Supplementary Table 3) requiring further evaluation as described below.

From the 863,904 Infinium EPIC CpG probes designed for the human genome, 402,935 (46.6%) mapped to the *Macaca mulatta* genome according to our parameters, from which 54,789 (13.6%) were perfect matches, 97,505 (24.2%) had 1 mismatch, 125,108 (31.0%) had 2 mismatches and 125,533 (31.2%) had 3 mismatches. Of these, 267,237 probes (66.3%, *versus* 74.3% in Infinium 450K) were identified as MM-functional and 135,698 were identified as MM-non-functional (Table 1). Of these MM-functional probes, 72,391 presented at least one mismatch at 10 bp or less from the CpG (Supplementary Table 3).

As above, we studied more carefully the distribution of all the substitutions from the whole mapped probes set (Supplementary Tables 6 and 7) and found a very similar distribution in *Macaca mulatta* as in *Chlorocebus sabaeus*.

Altogether, for their use in DNA methylation analysis using the two human BeadChips, 31.0 (EPIC) and 35.0 % (450K) of the probes could be classified as functional for *Macaca mulatta*.

These results showed in summary, that results were similar for the two species and allowing up to three mismatches for the 50 base pair probes potentially about a third of the probes on the respective arrays were targeting CpG positions in the two simian genomes. There was a strong overlap between the two simian species with 195,013 (about 85%) and 341,567 (about 84%) mapped probes in common between the two species and 139,085 (about 82%) and 216,272 (about 80%) probes in common for the Infinium 450K and Infinium EPIC microarrays, respectively.

As this probe set contained nonetheless probes which contained mismatches (outside the CpG position), which could have on influence on the probe behavior, a point which has been neglected in most studies so far. Therefore, final validation and refinement of the reliable probe sets required experimental analysis of samples from the respective species.

### Validation and refinement of our selection criteria for the Infinium 450K BeadChip

To determine to which extent CpG probes designed for the human genome and mapping to simian genomes could efficiently be used to detect methylation on these two simian models, we analyzed 25 CD4+ T cell samples from *Macaca mulatta* and 21 from *Chlorocebus sabaeus* on the Infinium 450K array (Figure 2). The density distribution of beta-values of the probes identified as perfectly matching were very similar to the bimodal beta-values density distribution commonly observed for human samples [40]. Beta-values density distribution of probes containing a single mismatch remained close to beta-values density distribution of perfectly matched probes, and this density distribution was still similar for probes with two mismatches. However, probes containing three mismatches presented density distribution of beta-values which started to deviate from the expected bimodal distribution and the distribution for unmapped probes did no longer follow the expected distribution comforting our restriction to a maximum of three mismatches to be present. These results were similar for both species.

**Figure 2:**
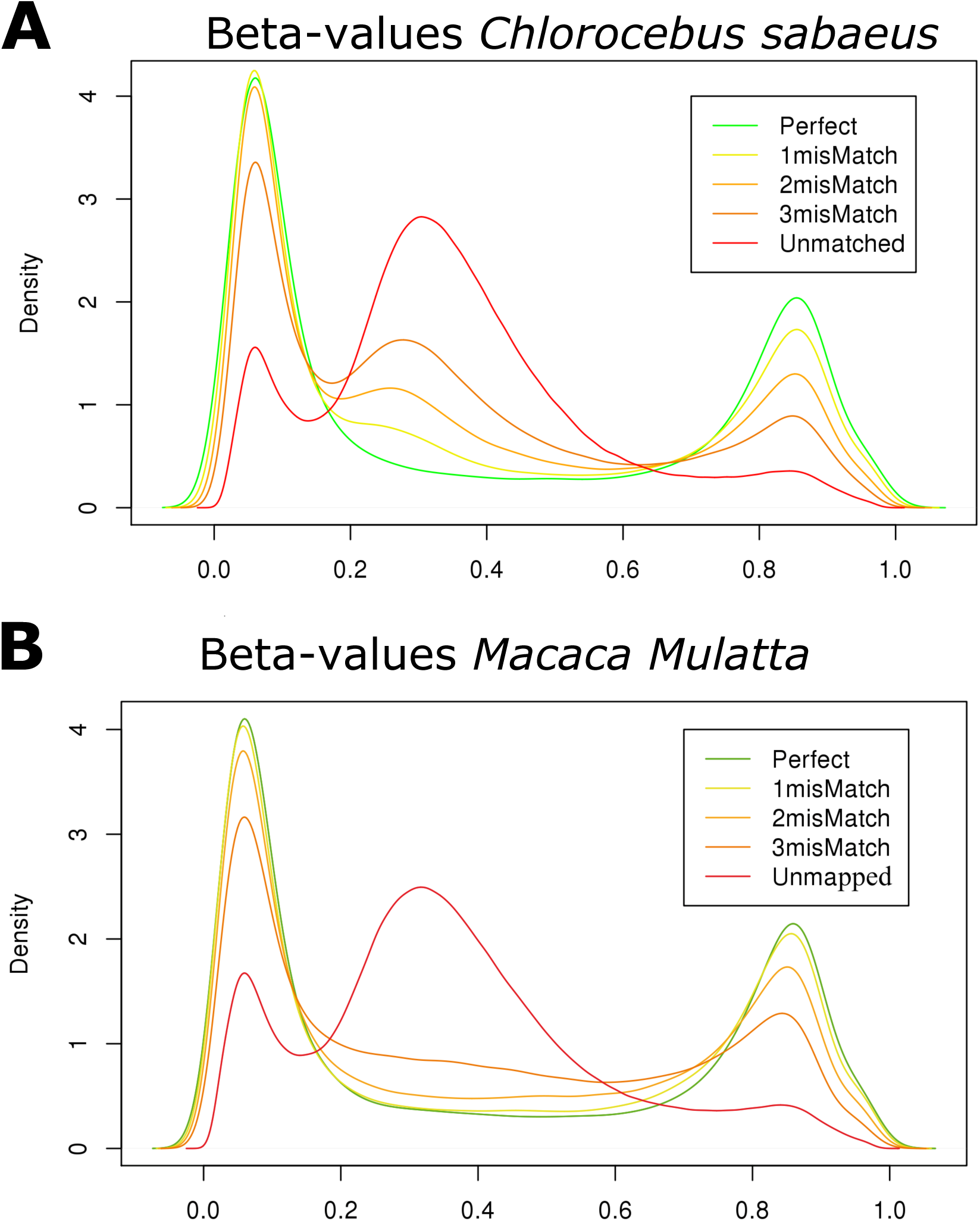
Beta-value density distribution for mapped and unmapped Infinium 450K CpG probes on *Chlorocebus sabaeus* (A) and *Macaca mulatta* (B) genome. Mapped probes are classified into categories representing probes with a perfect match, 1, 2 or 3 mismatches. For probes with 3 mismatches (dark orange) the beta-value distribution start to deviate from the bimodal distribution observed for probes with fewer or no mismatches. Unmapped probes (in red) have an aberrant beta-value distribution.

When analyzing the beta-values density distribution of probes in function of the position of the mismatch, we observed that, in both species, the beta-values distribution was closely related to the mismatch position (Figure 3). Thus, probes with a mismatch localized at 1 or 2 bp from CpG site presented an aberrant density distribution of beta-values among valid probes whereas probes containing a mismatch at 3-4 bp or more away from the CpG were similar to probes without mismatches independent of the number of mismatches present (one to three). We thus chose to remove probes presenting a mismatch at 1 or 2 bp from the CpG from the list of functional Infinium probes on simian models for both 450K and EPIC arrays. With this last filter, 162,053 CS-valid and 158,970 MM-valid probes remained for Infinium 450K (i.e. 33% of the Infinium 450K CpG probes), among which 130,484 (80.5% and 82.1%, respectively) were common to both species. For the Infinium EPIC array, 255,444 CS-valid and 252,235 MM-valid probes remained after filtering out probes presenting a mismatch at 1 or 2 bp from the CpG (i.e. 29% of the Infinium EPIC CpG probes). 202,495 (79.4% and 80.5%, respectively) of them were common to both species.

**Figure 3:**
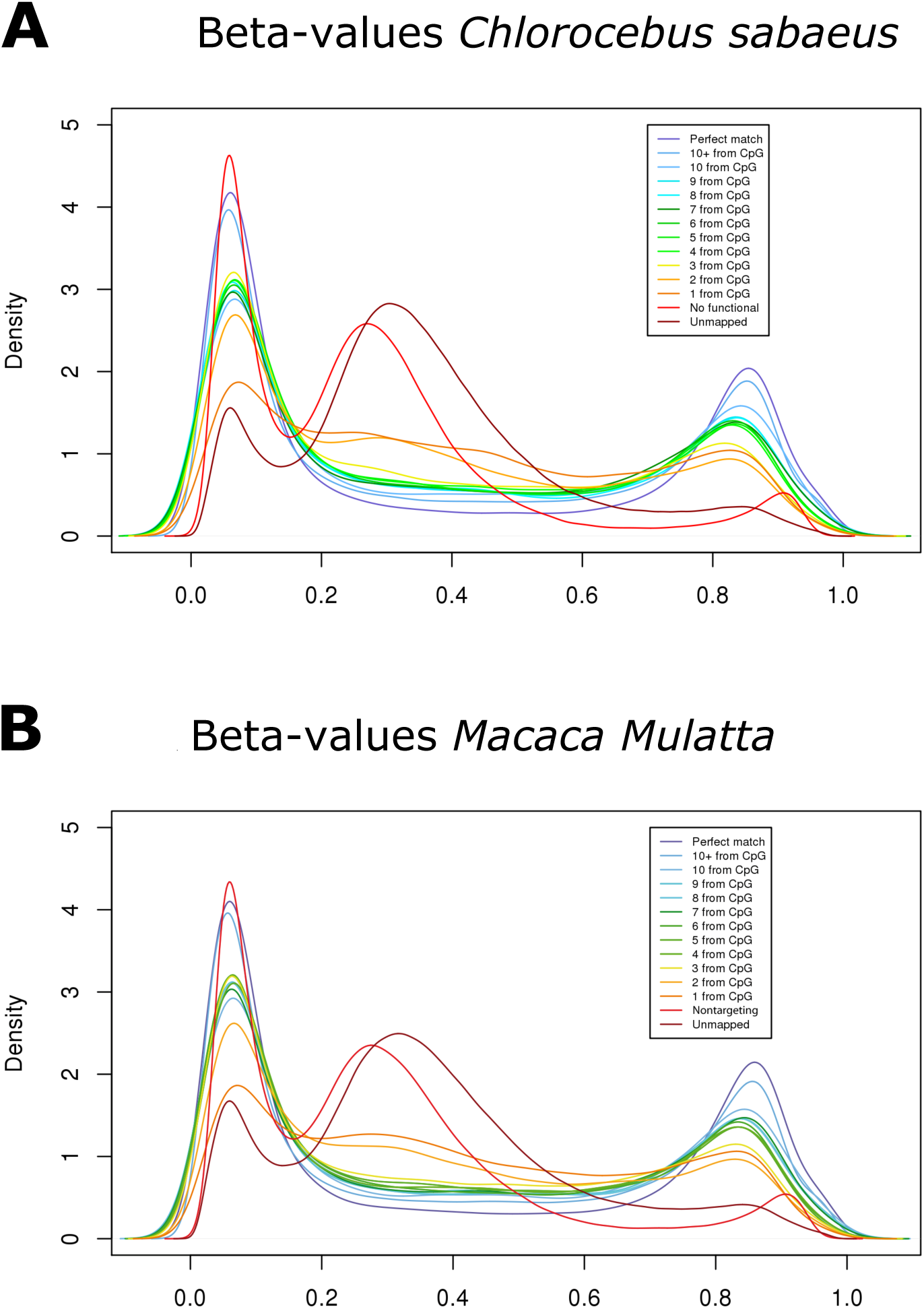
Beta-value density distribution for mapped and unmapped Infinium 450K CpG probes on *Chlorocebus sabaeus* (A) and *Macaca mulatta* (B) genome. Mapped probes are classified into categories representing probes with a perfect match and according to the mismatch position localized from 1 bp to 10 bp or more from CpG dinucleotide. Probes containing a mismatch localized at 1 or 2 bp (dark orange and orange) do no longer show the expected beta-value density distribution compared to the other mapped probes (yellow, greens, blues and purple). Non-targeting probes (red) and unmapped probes (dark red) have an aberrant beta-value distribution.

Furthermore, using Pyrosequencing, which as a sequencing-by-synthesis method is not dependent on human probes but uses species-specific amplification and sequencing primers, we validated DNA methylation levels measured by the respective Infinium probes at three CpG positions in each species showing a high correlation between the two orthogonal technologies and validating our approach of selecting reliable probes (Figure 4). The selected probes had a either or one mismatch (cg07181702, cg20733663, cg17245135 (n=0), cg21758672, cg09825979 cg15544721 (n=1)). There was no correlation between the presence and the position of the mismatch and the accuracy of the Infinium data compared to the Pyrosequencing data.

**Figure 4:**
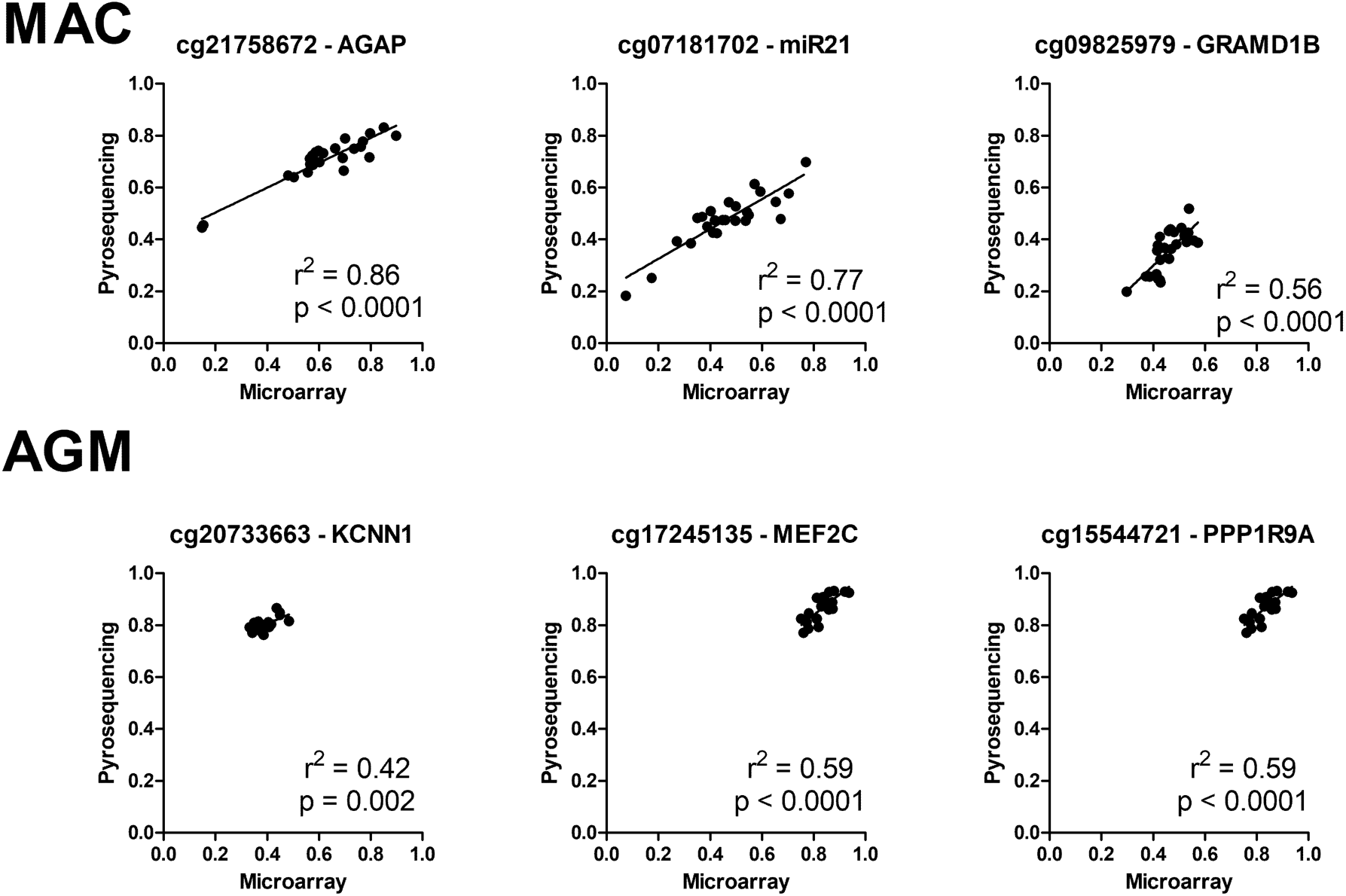
Validation of DNA methylation measures on CD4 T cells obtained on the Infinium 450K BeadChip by Pyrosequencing using locus-specific amplification primers designed for each species (Supplementary Table 1). Top row: *Macaca mulatta*, bottom row: *Chlorocebus sabaeus*.

### Annotation of valid probes

The annotation of the probes to the *Homo sapiens* genome (GRCh37) as described in the manifest files for both microarrays showed that the additional content of the Infinium EPIC BeadChip compared to the Infinium 450K array was mainly located in gene bodies, and in the open sea using the gene feature and CpG island feature annotation, respectively (Figure 5).To provide the user with similar information as contained in the Illumina manifest for the human genome, we annotated the location of the CpGs with matching probes according to the Ensembl gene annotation (version 84).

**Figure 5:**
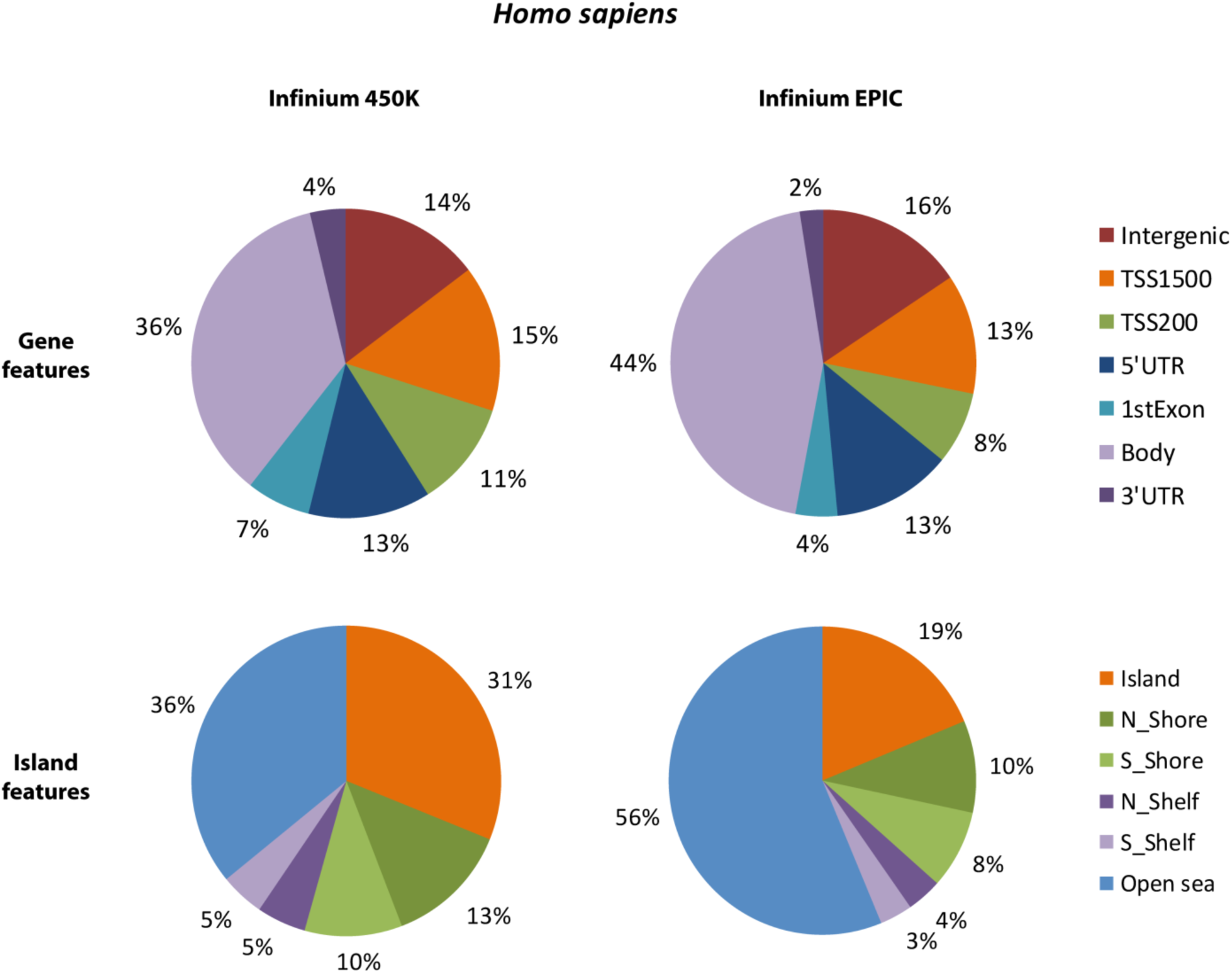
Distribution of human CpG probes in Infinium 450K and Infinium EPIC, as annotated by Illumina for *Homo sapiens* genome (GRCh37). All CpG probes are valid for human (482,421 probes in Infinium 450K and 863,904 in Infinium EPIC), which represent s 802,912 and 1,590,198 gene features annotations in total for Infinium 450K and Infinium EPIC, respectively. Note that one probe can be attributed to different transcripts and thus gene features, whereas one probe is attributed to only one island feature.

### Chlorocebus sabaeus

Among the 162,053 CS-valid probes on the Infinium 450K BeadChip, 102,674 were annotated for a gene feature (representing 16,259 genes) and 107,906 for a CpG island feature using the CS genome. 74,858 CS-valid probes were annotated for both a gene and an island feature. CS-valid probe sequences principally targeted CpGs located around transcription start sites and gene bodies (about 65% in total) and around 35% of the CS-valid probes target CpG in intergenic regions (*versus* 14% in human). According to UCSC islands prediction (see Materials and Methods section), more than 37% of human-designed CS-valid probes target CpG probes located in CpG islands, which is similar to the proportion for the human genome (Figure 6).

**Figure 6:**
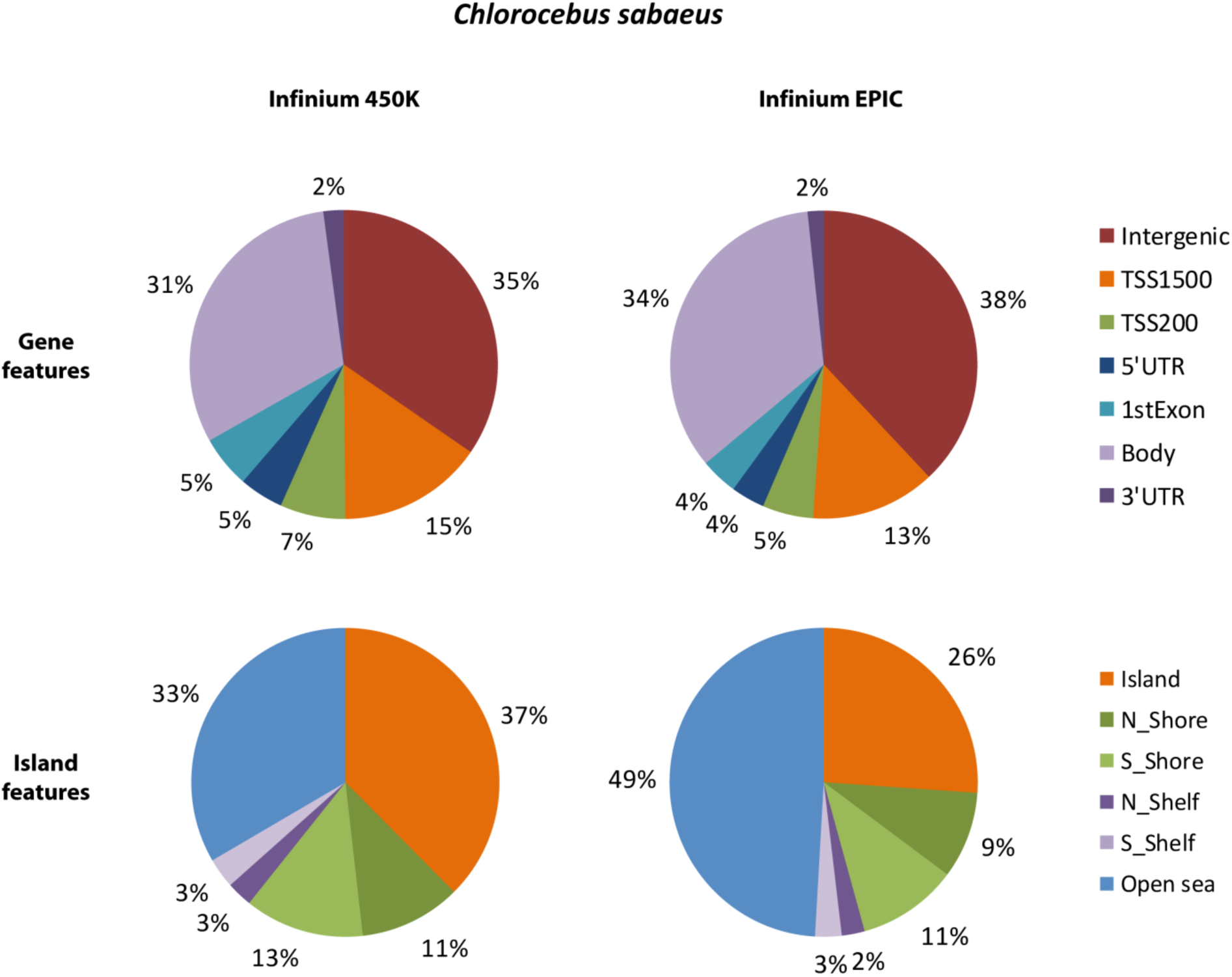
Distribution of human-designed CS-valid CpG probes in Infinium 450K and Infinium EPIC, as annotated for *Chlorocebus sabaeus* genome (Ensembl ChlSab1.1 version 84). We defined 162,053 CS-valid probes for Infinium 450K and 255,444 for Infinium EPIC, representing 171,608 and 268,062 gene feature annotations in total for Infinium 450K and Infinium EPIC, respectively. Note that one probe can be attributed to different transcripts and thus gene features, whereas one probe is attributed to only one island feature.

Among the 255,444 CS-valid probes on the Infinium EPIC BeadChip, 153,571 were annotated for a gene feature (representing 17,706 genes) and 130,041 for a CpG island feature. 90,169 probes were annotated for both a gene and an island feature. The proportion of the respective gene feature and CpG islands categories followed the trend observed for the human genome with an increased proportion in intergenic regions and gene bodies on the EPIC arrays (Figure 6).

### Macaca mulatta

Among the 158,970 MM-valid probes on the Infinium 450K array, 104,571 were annotated to a gene feature (representing 16,611 genes) and 98,884 to a CpG island feature (Figure 7). 73,352 MM-valid probes were annotated to both a gene and an island feature. As for CS-valid probes, but even more pronounced, MM-valid probe sequences principally targeted CpGs located in TSS regions and gene bodies (about 78% in total, 22% in intergenic regions). The distributions among island features according to the UCSC CpG island prediction was more similar between species (Figures 5, 6).

**Figure 7:**
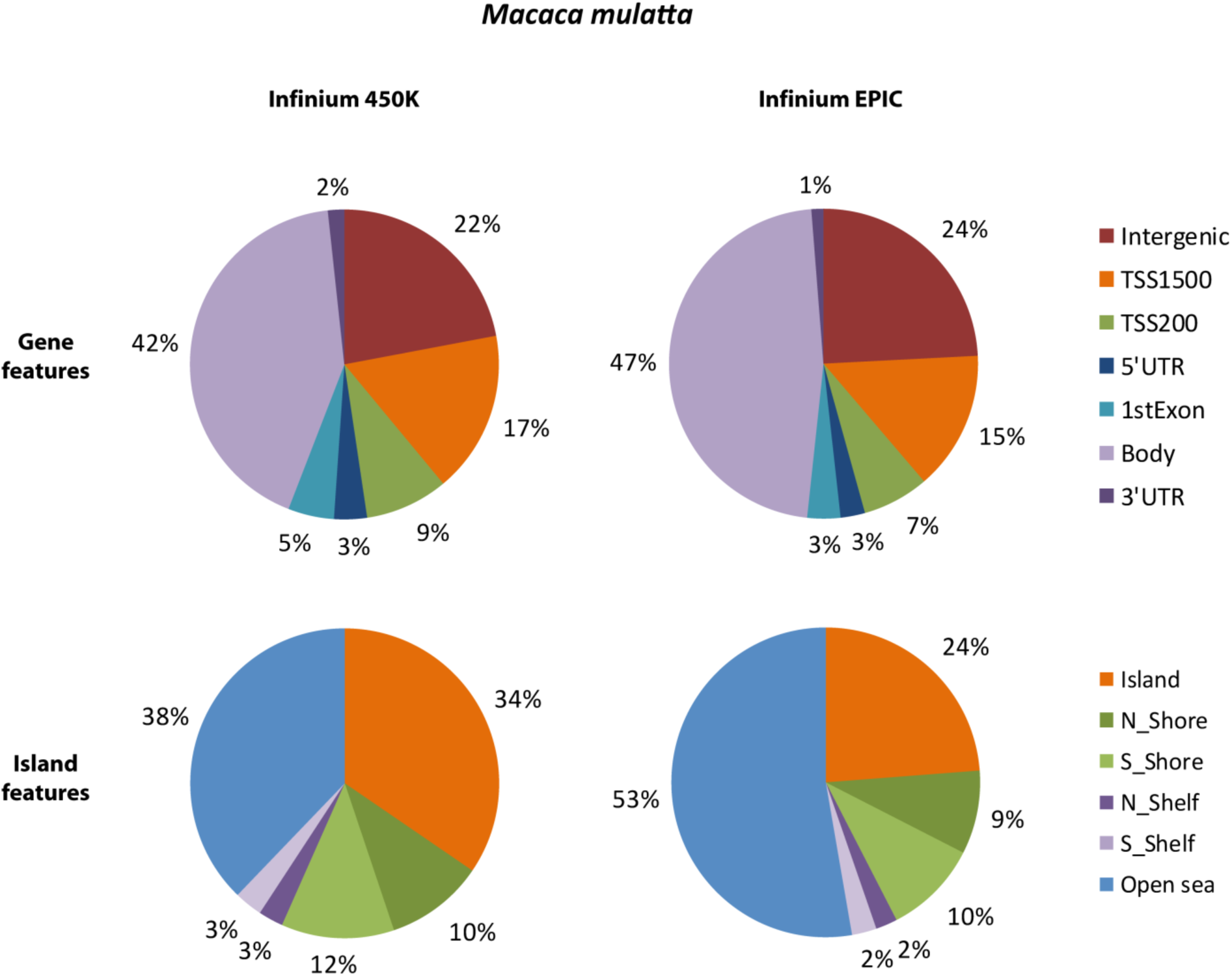
Distribution of human-designed MM-valid CpG probes in Infinium 450K and Infinium EPIC, as annotated for *Macaca mulatta* genome (Ensembl MMUL1.0 version 84). We defined 158,970 MM-valid probes for Infinium 450K and 252,235 for Infinium EPIC, representing 246,788 and 390,548 gene feature annotations in total for Infinium 450K and Infinium EPIC, respectively. Note that one probe can be attributed to different transcripts and thus gene features, whereas one probe is attributed to only one island feature.

Among the 252,235 MM-valid probes from Infinium EPIC mapped on the *Macaca mulatta* genome, 157,737 were annotated for a gene feature (representing 18,240 genes) and 119,248 for a CpG island feature. 88,355 MM-valid probes were annotated for both a gene and an island feature.

Compared to *Homo sapiens*, reliable Infinium 450K and EPIC probes followed the overall distribution of probes when using the CpG island feature annotation, while for the gene feature annotation the proportion of probes in intergenic regions was increased especially in *Chlorocebus sabaeus* at the expense of mainly gene body probes (Figures 5, 6 and 7). Overall 70 % of all simian genes were covered by at least one valid probe.

### 450K and EPIC manifest files for Chlorocebus sabaeus and Macaca mulatta

We provide the scientific community with new manifest files for Infinium 450K and EPIC BeadChips, adapted for genome-wide DNA methylation studies in *Chlorocebus sabaeus* or *Macaca mulatta* (Supplementary Material). We provide for each microarray and each species two files: one containing the whole set of CS-valid or MM-valid probes (filtered for probes with a mismatch at 1 or 2 bp from the CpG) and another file containing only perfectly matched probes. These annotation files retain the format of the original Illumina manifest and can thus be used without further modifications in analysis pipelines for BeadChips such as SQN [40] or the widely used ChAMP pipeline [42]. All columns of the respective manifest files are described in detail in the supporting file (manifest header descriptions file included in the compressed Supplementary Material). The last column details the number of mismatches as well as the string for the position of the mismatch returned by Bowtie. The interested user can thus select a subset of probes based on more stringent parameters, if needed. Of note, the string for the mismatch of the positions should be reversed for probe sequences mapping on the reverse strand of the monkey genomes.

## Discussion

Epigenome-wide association studies have recently been performed on many phenotypes, traits and diseases including cancer, immune, neurodegenerative and infectious diseases, with now more than 500 EWAS published [43] and many more large-scale studies are likely to be conducted in the near future, linking complex diseases and traits with changes in the epigenome. Furthermore, DNA methylation holds the promise to explain at least a part of the influences the environment has on a phenotype [44]. Cell lines or blood cells do in most cases not appropriately recapitulate the phenotype of complex diseases, requiring the use of tissue or animal models to further our understanding of disease etiology and evaluate potential future treatments. CS and MM represent reference models in biomedical research and due to the recently realized importance of epigenetic for human disease, it would be of importance to include DNA methylation analysis in comprehensive multi-level -omics analyses. However, few tools are currently available.

In the presented work we determined which probes of the most widely used DNA methylation arrays can be used to reliably analyze CpGs in the genomes CS and MM. The valid probes were very similar between *CS* and *MM*, with over 80% of mapped and valid CpG probe sequences in common between the two species. This is in agreement with the phylogenic proximity of these two *Cercopithecidae* [45]. However, the percentage of probes that can be reliably used differ from the one identified in a previous study conducted on *M. fascicularis* [19], a species closely related to *M. mulatta*, in which 61% of human probes mapped to the simian genome in contrast to the 47.6% identified here for *M. mulatta*. This difference can be explained by the more stringent parameters for mapping probe sequences in our study rather than differences between the two simian genomes. For instance, we allowed only three mismatches instead of the four mismatches allowed in the study by Ong et al. [19] to align Infinium 450K probe sequences on the simian genomes as we observed that beta-values densities showed aberrant distributions for probes with more than three mismatches as well as probes with fewer mismatches close to the targeted CpG. This feature has not been considered in previous studies, but had clearly strong influence on the measured DNA methylation levels (Figure 3). Furthermore, in contrast to Ong et al., [19] we only kept uniquely matched sequences on simian genomes because probes measuring DNA methylation levels at CpG in multiple genomic regions are difficult to interpret and thus are not informative as has been demonstrated in human studies of the arrays [46]. Furthermore, we have annotated 75.4% of aligned probes sequences (*versus* 94% in Ong et al). This can be explained because we only annotated probe sequences localized in genes or at less than 2,000 bp from a gene using a similar annotation format to the one provided by Illumina for the human genome. A similar approach was also proposed for the analysis of DNA methylation and, following oxidative bisulfite treatment, hydroxymethylation in MM, allowing up to four mismatches [18]. Again, position of potential mismatches was not further evaluated. The same filtering approach as proposed by Ong et al. [19] was subsequently also used for a study on osteoarthritis using baboon samples which concluded that a total of 44 % of probes on the 450K BeadChip could be reliably used [20].

Recently, human DNA methylation capture panels were adapted to the analysis of DNA methylation of non-human primates and notably AGM. While this approach allows to capture a larger proportion of regulatory elements in the simian genomes [47], it comes at a higher price and thus less well suited for projects with larger number of samples. Furthermore, in quantitative comparisons the Infinium BeadChips outperformed the sequencing-based approach in terms of quantitative precision even when pooling CpG levels of closely neighbored CpGs in the sequencing approach [48]. These results were obtained in human samples, it can thus be expected that the quantitative ratios will be increasingly distorted in the presence of sequence mismatches between the capture probes and the simian genomes.

*Homo sapiens* and *Macaca mulatta* have diverged about 25 million years ago and their sequence identity is about 93% [49]. At random, we should therefore expect about 86.5% of CpG sites in common between *Macaca mulatta*, and humans. Nevertheless, only 74.3% of mapped probes sequences were identified as MM-targeting. Analyzing more precisely the distribution of mismatches, we observed that C➔T and G➔A (Strong to weak, S➔W) nucleotide substitutions accounted for 28.8% of mismatches, in line with the fact that CpG sites are privileged mutational hot spots during speciation, due to methylation of cytosines [2–4]. Of note, W➔S nucleotide substitutions represented 38.3% of mismatches and around 60% when looking at CpG sites in agreement with the weak-to-strong bias observed during recent human evolution [50–52]. The same results were obtained for *Chlorocebus sabaeus* and similar results were found with the Infinium EPIC. For the latter, we observed a slightly increased substitution rate at CpG sites concomitant with a slightly lower number of valid probes compared to Infinium 450K for both species. This observation might be explained by the localization of the 350,000 additional probes in enhancers on the EPIC arrays as regulatory regions have been suggested to be fast evolving regions during human divergence from other primates [53,54]. As the overall percentage of reliable probes between the different generations of BeadChips remains similar, it can be expected that future human generations of human Infinium DNA methylation microarrays, will also improve the coverage of the simian genomes.

Annotation of these valid probes for gene features showed higher proportion of probes targeting CpGs located in intergenic regions compared to CpGs in promoter or gene regions. This tendency was increased in *Chlorocebus sabaeus* compared to *Macaca mulatta.* In contrast, the frequency of location of valid probes within CpG island features were similar between the two species. It is important to point out the different annotation levels between these distinct species, which impacts the number of identified/predicted genes and transcripts in each species. Indeed, human GRCh37 annotation (version 75 from Ensembl) contains 57,952 genes and pseudogenes for 196,501 transcripts plus thousands of alternate sequences, whereas 30.246 genes are annotated in *Macaca mulatta* for 44.725 transcripts and 27,985 genes are annotated in *Chlorocebus sabaeus* for only 28,078 transcripts [55,56]. CpG island feature annotations, in counterpart, are directly predicted from reference sequence. This might explain the observed differences between the species depending on the genomic region. It is indeed important to keep in mind that simian annotations generally are still less precise and documented than human annotation and this lack of information may contribute to the observed differences.

Nonetheless, we provide to the research community a list of 162,053 annotated CS-valid probes corresponding to 16,259 genes for *Chlorocebus sabaeus* and a list of 158,970 annotated MM-valid probes corresponding to 16,611 genes for *Macaca mulatta* for Infinium 450K array, as well as a list of 255,444 annotated CS-valid probes corresponding to 17,706 genes for *Chlorocebus sabaeus* and a list of 252,235 annotated MM-valid probes corresponding to 18,240 genes for *Macaca mulatta* for the newly released Infinium EPIC array. While previous studies identified list of probes, they did not provide a species-specific annotation file. Furthermore, we annotated probes according to Illumina gene and CpG island features and built annotation files following the Illumina manifest file format. We kept the human GRCh37 annotation for comparison purpose and our annotation files can directly be used with a number of processing tools. Although the proportions of the different gene features are altered compared to the distribution for the human genome with a reduction in gene bodies and around the TSS, the current generation of the methylation microarray cover around 60 % of all known genes in the two species and these genes are covered by two to six probes allowing multiple reliable measurements of DNA methylation to increase confidence in the obtained data. This observation will also be reinforced by the fact that human array contains proportionally more probes in the promoter regions (often associated with CpG islands) ensuring that promoter CpGs regions remain sufficiently covered. While genes of special interest for a research group might be missing from these lists, the arrays provide the currently most comprehensive tool for DNA methylation analysis at a reasonable price. The number of available reliable probes exceeds by several orders of magnitude the number of probes available on the first generation of the Infinium DNA methylation BeadChips, the 27K array, which led to the discovery of DNA methylation changes following environmental exposure such as tobacco smoking [57] or disease associated changes [58,59], and whose success led to the development of the higher-density DNA methylation arrays evaluated in this study. While monkey-specific microarrays would constitute ideal tools, however, commercial of-the-shelf DNA methylation microarrays for model organisms have been awaited in vain. On the other hand, comprehensive whole genome bisulfite sequencing projects remain for most laboratories prohibitively expensive when performed at coverage allowing reliable methylation calls.

In summary, our approach using standard bioinformatic tools and validated quality criteria using orthogonal analysis technology concerning notably the importance of the number and position of any mismatches should be applicable to other simian and perhaps even other mammalian species. Our approach allows a rapid selection of reliable probes for any organism with an annotated genome sequence. Of course, a more divergent genome sequence, will lead to a fewer probes that can be used with high confidence and for some organisms this might eventually lead to an unfavorable cost / output balance, such as mice where only 10-12k probes for the 450K BeadChip and 13K for the EPIC array were found reliable [18,60,61].

In conclusion, the presented work investigated in depth the suitability of human DNA methylation Infinium BeadChips for the use in two widely used simian model species. The annotation files are provided in a format compatible with a variety of preprocessing, normalization and analytical pipelines designed for data analysis from 450K/EPIC arrays facilitating high-throughput DNA methylation analyses in *Macaca mulatta* and *Chlorocebus sabaeus* for many questions of biomedical relevance.

## Supporting information

Supplementary Material_manifest files

## Acknowledgements

We would like to acknowledge grant support from ANRS and the institutional budget form the CNRGH. SP was recipient of a PhD fellowship from the University Paris Diderot, Sorbonne Paris Cité.

## Notes

### Competing Interest Statement

The authors have declared no competing interest.

