## Supplementary Tables And Figures for "Analysis and annotation of genome-wide DNA methylation patterns in two nonhuman primate species using the Infinium Human Methylation 450K and EPIC BeadChips"

| Species | Gene | ProbeID | Coordinates | Size | Forward Primer | Reverse Primer | Polymerase | Tp (°C) | Pyrosequencing Primer | CpG |
|---|---|---|---|---|---|---|---|---|---|---|
| **AGM** | *KCNN1* | cg20733663 | chr6:16469411-16470348 | 338 | Biotine-GTTTGGATGYGTTGGGTGTT | TCCAACCCTACCCTCTACCC | Qiagen | 66.5 | CAAAATTTAAAAATCTAATA | 7-1 |
| | *MEF2C* | cg17245135 | chr4:82706770-82706907 | 138 | GTTTGATTGAGGGATTTTTTTT | Biotin-AATCTCCTATTAACAACTTAAACAACTATA | Phusion | 58.8 | TGAGTTTAATGGGGGA | 1-7 |
| | *PPP1R9A* | cg15544721 | chr21:54129632-54129824 | 193 | TTGGGGTTTTATTTAGTTTGAGTTT | Biotin-CCTCATCCCTTTCTCTTTAACAATA | Phusion | 61.5 | AAATTTTAGGTTATTTTGT | 1-2 |
| **MAC** | *AGAP* | cg21758672 | chr12:99801709-99801929 | 221 | AAAGATTTAGTATTAGGTTTTTTTTGTAGA | Biotin-ATCTACTCCAAATCACAAACTTTTC | Phusion | 53.8 | ATTAGTTGTTGTAGTAGGA | 1-3 |
| | | | | | | | | | TTTTAAATTGAGTTAGGAGA | 4-8 |
| | *MIR21* | cg07181702 | chr16:44086577-44086773 | 197 | TTGATGTTGATTGTTGAATTTTATG | Biotine-TCCCTCCACACTACTACATTATAACAC | Qiagen | 61.5 | AATTTTATGGTAATATTAGT | 1 |
| | *GRAMD1B* | cg09825979 | chr3:99085974-99086191 | 218 | Biotin – TTTTTAGGTTTAATTTTAATTGTTGTATAA | TACCAAATTCCCACACACAAAC | Phusion | 56.1 | AAAAATCATCTTTTTACA | 6-1 |

**Supplementary Table 1**: Primers and PCR conditions used for Pyrosequencing-based validation of DNA methylation levels.

| | *Chlorocebus sabaeus* | | *Macaca mulatta* | |
|---|---|---|---|---|
| | **238383** | **100.00%** | **233589** | **100.00%** |
| Substitution in C | 30353 | 12.7% | 29599 | 12.7% |
| A | 17780 | 7.5% | 4026 | 1.7% |
| G | 17959 | 7.5% | 4565 | 2.0% |
| T | 26338 | 11.0% | 21008 | 9.0% |
| Substitution in G | 31702 | 13.3% | 24983 | 10.7% |
| A | 26901 | 11.3% | 21166 | 9.1% |
| C | 18478 | 7.8% | 4513 | 1.9% |
| T | 18047 | 7.6% | 3950 | 1.7% |

**Supplementary Table 2:** Repartition of substitutions occurring at CpG sites in non-targeting Infinium 450K microarray 50 bp probe sequences mapped on simian genomes.

| Mismatch | Infinium 450K | | Infinium EPIC | |
|---|---|---|---|---|
| n# bases from CpG | *Chlorocebus sabaeus* | *Macaca mulatta* | *Chlorocebus sabaeus* | *Macaca mulatta* |
| 1 | 4694 | 4505 | 7436 | 7235 |
| 2 | 5165 | 4991 | 8042 | 7767 |
| 3 | 4889 | 4862 | 7808 | 7764 |
| 4 | 5092 | 4885 | 8017 | 7787 |
| 5 | 4799 | 4675 | 7546 | 7389 |
| 6 | 4619 | 4602 | 7411 | 7310 |
| 7 | 4435 | 4413 | 7280 | 7134 |
| 8 | 4411 | 4349 | 7019 | 6906 |
| 9 | 4170 | 4128 | 6697 | 6639 |
| 10 | 4183 | 4052 | 6597 | 6460 |

**Supplementary Table 3:** Repartition of the number of probes according to the distance of the closest mismatch to the CpG site on the Infinium 450K and EPIC microarray.

| 450K | *Chlorocebus sabaeus* | | | | | | | | | | | | |
|---|---|---|---|---|---|---|---|---|---|---|---|---|---|
| **Base** | **A->C** | **A->G** | **A->T** | **C->A** | **C->G** | **C->T** | **G->A** | **G->C** | **G->T** | **T->A** | **T->C** | **T->G** | **ALL** | ***Percent** |
| **1** | 2322 | 682 | 114 | 212 | 282 | 606 | 664 | 2605 | 214 | 124 | 11661 | 187 | 19673 | 8.5% |
| **2** | 173 | 11093 | 113 | 178 | 2414 | 599 | 641 | 205 | 198 | 140 | 600 | 2078 | 18432 | 8.0% |
| **3** | 283 | 1218 | 182 | 320 | 428 | 1214 | 1032 | 343 | 338 | 186 | 1055 | 290 | 6889 | 3.0% |
| **4** | 287 | 1172 | 194 | 330 | 392 | 1216 | 1213 | 410 | 366 | 195 | 1148 | 339 | 7262 | 3.1% |
| **5** | 257 | 1108 | 172 | 361 | 373 | 1151 | 1146 | 423 | 359 | 180 | 1079 | 293 | 6902 | 3.0% |
| **6** | 280 | 1185 | 173 | 325 | 402 | 1208 | 1214 | 407 | 332 | 171 | 1110 | 269 | 7076 | 3.1% |
| **7** | 260 | 1207 | 191 | 347 | 385 | 1246 | 1175 | 433 | 360 | 195 | 1116 | 290 | 7205 | 3.1% |
| **8** | 308 | 1154 | 181 | 361 | 419 | 1148 | 1276 | 430 | 362 | 175 | 1150 | 305 | 7269 | 3.1% |
| **9** | 303 | 1155 | 173 | 311 | 366 | 1210 | 1192 | 382 | 322 | 173 | 1111 | 254 | 6952 | 3.0% |
| **10** | 284 | 1192 | 156 | 391 | 415 | 1202 | 1213 | 402 | 319 | 171 | 1186 | 268 | 7199 | 3.1% |
| **11** | 241 | 1093 | 165 | 318 | 368 | 1169 | 1104 | 407 | 348 | 150 | 1123 | 275 | 6761 | 2.9% |
| **12** | 258 | 1122 | 190 | 322 | 364 | 1161 | 1224 | 413 | 328 | 187 | 1171 | 298 | 7038 | 3.0% |
| **13** | 281 | 1182 | 146 | 315 | 364 | 1179 | 1180 | 379 | 313 | 192 | 1188 | 249 | 6968 | 3.0% |
| **14** | 239 | 1220 | 156 | 350 | 377 | 1152 | 1228 | 371 | 344 | 141 | 1197 | 258 | 7033 | 3.0% |
| **15** | 269 | 1143 | 155 | 333 | 391 | 1202 | 1214 | 384 | 322 | 178 | 1157 | 256 | 7004 | 3.0% |
| **16** | 284 | 1171 | 163 | 300 | 409 | 1164 | 1165 | 409 | 329 | 187 | 1135 | 269 | 6985 | 3.0% |
| **17** | 262 | 1159 | 201 | 317 | 416 | 1179 | 1216 | 374 | 308 | 158 | 1166 | 265 | 7021 | 3.0% |
| **18** | 282 | 1149 | 188 | 337 | 382 | 1206 | 1168 | 400 | 349 | 174 | 1162 | 268 | 7065 | 3.1% |
| **19** | 298 | 1140 | 162 | 327 | 407 | 1231 | 1120 | 378 | 313 | 168 | 1193 | 256 | 6993 | 3.0% |
| **20** | 235 | 1186 | 164 | 309 | 383 | 1170 | 1153 | 397 | 344 | 155 | 1145 | 278 | 6919 | 3.0% |
| **21** | 260 | 1162 | 159 | 332 | 390 | 1194 | 1117 | 418 | 332 | 174 | 1173 | 281 | 6992 | 3.0% |
| **22** | 284 | 1103 | 169 | 325 | 356 | 1231 | 1097 | 390 | 334 | 193 | 1180 | 272 | 6934 | 3.0% |
| **23** | 239 | 1235 | 174 | 353 | 429 | 1159 | 1138 | 394 | 359 | 164 | 1172 | 279 | 7095 | 3.1% |
| **24** | 281 | 1189 | 173 | 303 | 401 | 1204 | 1201 | 358 | 329 | 166 | 1131 | 265 | 7001 | 3.0% |
| **25** | 285 | 1118 | 185 | 318 | 364 | 1192 | 1144 | 404 | 306 | 178 | 1194 | 279 | 6967 | 3.0% |
| **26** | 257 | 1182 | 193 | 307 | 380 | 1174 | 1205 | 401 | 336 | 189 | 1055 | 264 | 6943 | 3.0% |

| | | | | | | | | | | | | | |
|---|---|---|---|---|---|---|---|---|---|---|---|---|---|
| **27** | 261 | 1163 | 192 | 333 | 392 | 1152 | 1192 | 389 | 314 | 184 | 1219 | 272 | 7063 | 3.1% |
| **28** | 250 | 1178 | 155 | 329 | 406 | 1217 | 1188 | 398 | 310 | 187 | 1182 | 261 | 7061 | 3.1% |
| **29** | 281 | 1177 | 163 | 331 | 401 | 1166 | 1201 | 376 | 332 | 163 | 1172 | 260 | 7023 | 3.0% |
| **30** | 270 | 1160 | 159 | 364 | 401 | 1132 | 1150 | 426 | 328 | 167 | 1166 | 290 | 7013 | 3.0% |
| **31** | 264 | 1180 | 166 | 336 | 426 | 1147 | 1232 | 391 | 321 | 155 | 1211 | 270 | 7099 | 3.1% |
| **32** | 242 | 1272 | 150 | 326 | 397 | 1164 | 1218 | 378 | 334 | 177 | 1098 | 255 | 7011 | 3.0% |
| **33** | 292 | 1181 | 169 | 340 | 417 | 1234 | 1143 | 414 | 298 | 172 | 1118 | 270 | 7048 | 3.0% |
| **34** | 262 | 1167 | 190 | 321 | 407 | 1164 | 1140 | 378 | 331 | 169 | 1088 | 261 | 6878 | 3.0% |
| **35** | 274 | 1166 | 183 | 330 | 401 | 1145 | 1197 | 412 | 335 | 168 | 1126 | 260 | 6997 | 3.0% |
| **36** | 278 | 1201 | 177 | 320 | 347 | 1217 | 1216 | 369 | 333 | 195 | 1136 | 244 | 7033 | 3.0% |
| **37** | 277 | 1124 | 199 | 349 | 398 | 1209 | 1187 | 401 | 306 | 169 | 1138 | 300 | 7057 | 3.1% |
| **38** | 279 | 1167 | 173 | 289 | 400 | 1189 | 1206 | 361 | 332 | 180 | 1158 | 288 | 7022 | 3.0% |
| **39** | 277 | 1204 | 175 | 323 | 395 | 1170 | 1235 | 380 | 335 | 172 | 1062 | 255 | 6983 | 3.0% |
| **40** | 258 | 1148 | 180 | 349 | 389 | 1204 | 1187 | 389 | 358 | 151 | 1097 | 253 | 6963 | 3.0% |
| **41** | 288 | 1187 | 160 | 323 | 382 | 1082 | 1168 | 410 | 314 | 180 | 1138 | 287 | 6919 | 3.0% |
| **42** | 278 | 1178 | 160 | 323 | 392 | 1170 | 1168 | 385 | 306 | 171 | 1117 | 283 | 6931 | 3.0% |
| **43** | 246 | 1159 | 157 | 348 | 392 | 1226 | 1158 | 400 | 316 | 149 | 1149 | 265 | 6965 | 3.0% |
| **44** | 269 | 1203 | 149 | 343 | 407 | 1180 | 1179 | 373 | 332 | 172 | 1161 | 270 | 7038 | 3.0% |
| **45** | 251 | 1154 | 167 | 338 | 395 | 1232 | 1184 | 396 | 356 | 166 | 1206 | 286 | 7131 | 3.1% |
| **46** | 257 | 1107 | 179 | 370 | 391 | 1246 | 1222 | 375 | 333 | 194 | 1184 | 271 | 7129 | 3.1% |
| **47** | 293 | 1178 | 202 | 371 | 390 | 1152 | 1224 | 402 | 352 | 187 | 1212 | 251 | 7214 | 3.1% |
| **48** | 317 | 1068 | 191 | 335 | 390 | 1119 | 1160 | 466 | 346 | 197 | 1219 | 279 | 7087 | 3.1% |
| **49** | 2174 | 625 | 100 | 207 | 172 | 599 | 632 | 2448 | 186 | 98 | 11214 | 139 | 18594 | 8.0% |
| **50** | 201 | 11482 | 120 | 206 | 2775 | 601 | 659 | 270 | 204 | 110 | 681 | 2306 | 19615 | 8.5% |
| **Total** | 4.3% | 19.4% | 2.1% | 4.0% | 5.9% | 14.2% | 14.2% | 5.9% | 4.0% | 2.1% | 19.3% | 4.3% | 399452 | 231217 |

**Supplementary Table 4:** Repartition of substitutions along the 50 bp probe sequence for mapped Infinium 450K probes on the *Chlorocebus sabaeus* genome. *Percentages are calculated over the total number of mapped probes (231,217), NOT the total number of substitutions (399,452) as one probe can have several mismatches.

| EPIC | *Chlorocebus sabaeus* | | | | | | | | | | | | | |
|---|---|---|---|---|---|---|---|---|---|---|---|---|---|---|
| **Base** | **A->C** | **A->G** | **A->T** | **C->A** | **C->G** | **C->T** | **G->A** | **G->C** | **G->T** | **T->A** | **T->C** | **T->G** | **ALL** | ***Percent** |
| **1** | 5029 | 1149 | 222 | 356 | 437 | 1152 | 1156 | 5563 | 382 | 261 | 28467 | 314 | 44488 | 10.9% |
| **2** | 291 | 25902 | 222 | 317 | 4539 | 1086 | 1094 | 328 | 373 | 245 | 1028 | 4362 | 39787 | 9.7% |
| **3** | 459 | 2223 | 336 | 525 | 708 | 2120 | 1918 | 602 | 575 | 337 | 1949 | 502 | 12254 | 3.0% |
| **4** | 463 | 2079 | 347 | 584 | 618 | 2147 | 2148 | 635 | 621 | 331 | 2055 | 550 | 12578 | 3.1% |
| **5** | 463 | 2056 | 366 | 611 | 609 | 2043 | 2103 | 649 | 607 | 365 | 1959 | 514 | 12345 | 3.0% |
| **6** | 449 | 2096 | 340 | 554 | 641 | 2113 | 2168 | 660 | 596 | 322 | 2045 | 473 | 12457 | 3.0% |
| **7** | 460 | 2124 | 361 | 581 | 640 | 2250 | 2087 | 686 | 616 | 380 | 2014 | 478 | 12677 | 3.1% |
| **8** | 477 | 2040 | 338 | 593 | 647 | 2102 | 2192 | 681 | 609 | 310 | 2005 | 502 | 12496 | 3.1% |
| **9** | 466 | 2083 | 342 | 563 | 619 | 2183 | 2160 | 634 | 560 | 341 | 2024 | 430 | 12405 | 3.0% |
| **10** | 495 | 2116 | 326 | 642 | 642 | 2164 | 2111 | 641 | 549 | 320 | 2101 | 457 | 12564 | 3.1% |
| **11** | 422 | 2008 | 337 | 590 | 647 | 2055 | 2104 | 665 | 596 | 293 | 2020 | 482 | 12219 | 3.0% |
| **12** | 476 | 2009 | 339 | 590 | 608 | 2044 | 2141 | 659 | 602 | 334 | 2031 | 500 | 12333 | 3.0% |
| **13** | 489 | 2034 | 314 | 568 | 591 | 2133 | 2139 | 627 | 574 | 337 | 2091 | 455 | 12352 | 3.0% |
| **14** | 447 | 2120 | 312 | 571 | 598 | 2110 | 2166 | 611 | 605 | 292 | 2046 | 428 | 12306 | 3.0% |
| **15** | 483 | 2069 | 331 | 611 | 638 | 2119 | 2195 | 638 | 520 | 313 | 2072 | 465 | 12454 | 3.0% |
| **16** | 489 | 2128 | 303 | 524 | 668 | 2098 | 2141 | 646 | 567 | 339 | 2088 | 442 | 12433 | 3.0% |
| **17** | 436 | 2026 | 376 | 565 | 636 | 2124 | 2090 | 597 | 557 | 318 | 2084 | 455 | 12264 | 3.0% |
| **18** | 471 | 2148 | 325 | 584 | 656 | 2170 | 2134 | 627 | 575 | 318 | 2061 | 431 | 12500 | 3.1% |
| **19** | 498 | 2015 | 333 | 581 | 658 | 2128 | 2079 | 631 | 572 | 342 | 2052 | 430 | 12319 | 3.0% |
| **20** | 435 | 2084 | 331 | 533 | 616 | 2124 | 2079 | 615 | 576 | 279 | 2078 | 460 | 12210 | 3.0% |
| **21** | 450 | 2093 | 327 | 602 | 637 | 2153 | 2040 | 648 | 555 | 308 | 2103 | 464 | 12380 | 3.0% |
| **22** | 460 | 2045 | 337 | 543 | 610 | 2167 | 1982 | 623 | 572 | 343 | 2056 | 470 | 12208 | 3.0% |
| **23** | 417 | 2165 | 320 | 604 | 652 | 2107 | 2172 | 636 | 621 | 318 | 2096 | 482 | 12590 | 3.1% |
| **24** | 476 | 2107 | 338 | 524 | 627 | 2192 | 2125 | 593 | 576 | 326 | 2071 | 454 | 12409 | 3.0% |
| **25** | 479 | 2028 | 320 | 559 | 629 | 2169 | 2094 | 637 | 550 | 351 | 2065 | 466 | 12347 | 3.0% |
| **26** | 424 | 2075 | 345 | 528 | 647 | 2090 | 2169 | 661 | 574 | 334 | 1947 | 446 | 12240 | 3.0% |

| | | | | | | | | | | | | | |
|---|---|---|---|---|---|---|---|---|---|---|---|---|---|
| **27** | 451 | 2061 | 330 | 586 | 638 | 2080 | 2125 | 639 | 523 | 338 | 2135 | 435 | 12341 | 3.0% |
| **28** | 459 | 2125 | 310 | 558 | 644 | 2139 | 2112 | 627 | 538 | 339 | 2113 | 463 | 12427 | 3.0% |
| **29** | 490 | 2028 | 322 | 578 | 677 | 2060 | 2144 | 608 | 570 | 351 | 2069 | 460 | 12357 | 3.0% |
| **30** | 443 | 2074 | 325 | 579 | 639 | 2082 | 2039 | 675 | 553 | 300 | 2090 | 499 | 12298 | 3.0% |
| **31** | 491 | 2114 | 322 | 603 | 639 | 2064 | 2162 | 638 | 569 | 304 | 2119 | 469 | 12494 | 3.1% |
| **32** | 441 | 2183 | 296 | 579 | 625 | 2098 | 2250 | 609 | 565 | 330 | 2048 | 461 | 12485 | 3.1% |
| **33** | 480 | 2087 | 333 | 589 | 635 | 2183 | 2053 | 646 | 539 | 295 | 2048 | 448 | 12336 | 3.0% |
| **34** | 459 | 2054 | 343 | 555 | 643 | 2102 | 2086 | 602 | 533 | 315 | 2021 | 455 | 12168 | 3.0% |
| **35** | 484 | 2042 | 328 | 587 | 641 | 2080 | 2130 | 637 | 605 | 316 | 2008 | 429 | 12287 | 3.0% |
| **36** | 463 | 2110 | 323 | 555 | 608 | 2130 | 2114 | 621 | 556 | 342 | 1983 | 446 | 12251 | 3.0% |
| **37** | 477 | 1956 | 354 | 560 | 641 | 2161 | 2048 | 644 | 563 | 328 | 2051 | 455 | 12238 | 3.0% |
| **38** | 458 | 2087 | 321 | 537 | 651 | 2092 | 2165 | 587 | 575 | 342 | 2066 | 473 | 12354 | 3.0% |
| **39** | 482 | 2103 | 328 | 573 | 620 | 2114 | 2178 | 622 | 572 | 313 | 2020 | 453 | 12378 | 3.0% |
| **40** | 456 | 2090 | 343 | 585 | 648 | 2215 | 2105 | 617 | 600 | 296 | 1951 | 425 | 12331 | 3.0% |
| **41** | 467 | 2114 | 344 | 581 | 620 | 1970 | 2113 | 641 | 589 | 310 | 2021 | 476 | 12246 | 3.0% |
| **42** | 487 | 2122 | 320 | 557 | 631 | 2099 | 2131 | 637 | 577 | 342 | 2069 | 477 | 12449 | 3.0% |
| **43** | 435 | 2010 | 315 | 615 | 637 | 2173 | 2130 | 654 | 562 | 317 | 2055 | 448 | 12351 | 3.0% |
| **44** | 479 | 2146 | 302 | 591 | 646 | 2134 | 2162 | 583 | 570 | 313 | 2025 | 475 | 12426 | 3.0% |
| **45** | 417 | 2012 | 326 | 615 | 647 | 2174 | 2094 | 650 | 567 | 324 | 2133 | 482 | 12441 | 3.0% |
| **46** | 454 | 1984 | 301 | 626 | 602 | 2243 | 2191 | 598 | 553 | 362 | 2067 | 467 | 12448 | 3.0% |
| **47** | 490 | 2098 | 346 | 596 | 614 | 2045 | 2171 | 630 | 573 | 344 | 2148 | 449 | 12504 | 3.1% |
| **48** | 539 | 2003 | 328 | 542 | 630 | 2001 | 2089 | 732 | 546 | 333 | 2203 | 453 | 12399 | 3.0% |
| **49** | 4528 | 1123 | 194 | 352 | 325 | 1050 | 1176 | 4541 | 329 | 196 | 26147 | 240 | 40201 | 9.8% |
| **50** | 344 | 28179 | 236 | 362 | 5747 | 1102 | 1202 | 398 | 350 | 218 | 1178 | 4996 | 44312 | 10.8% |
| **Total** | 4.3% | 20.6% | 2.2% | 3.8% | 5.5% | 13.8% | 13.8% | 5.4% | 3.8% | 2.2% | 20.5% | 4.2% | 738137 | 408933 |

**Supplementary Table 5:** Repartition of substitutions along the 50 bp probe sequence for mapped Infinium EPIC probes on the *Chlorocebus sabaeus* genome. *Percentages are calculated over the total number of mapped probes (408,933), NOT the total number of substitutions (738,137) as one probe can have several mismatches.

| 450K | *Macaca mulatta* | | | | | | | | | | | | | |
|---|---|---|---|---|---|---|---|---|---|---|---|---|---|---|
| **Base** | **A->C** | **A->G** | **A->T** | **C->A** | **C->G** | **C->T** | **G->A** | **G->C** | **G->T** | **T->A** | **T->C** | **T->G** | **ALL** | ***Percent** |
| **1** | 2267 | 675 | 114 | 186 | 270 | 610 | 644 | 2660 | 210 | 121 | 11157 | 195 | 19109 | 8.4% |
| **2** | 169 | 10882 | 116 | 174 | 2375 | 608 | 627 | 188 | 182 | 114 | 583 | 2023 | 18041 | 8.0% |
| **3** | 236 | 1199 | 185 | 303 | 444 | 1203 | 1042 | 321 | 350 | 169 | 1105 | 274 | 6831 | 3.0% |
| **4** | 269 | 1148 | 196 | 312 | 374 | 1168 | 1161 | 396 | 354 | 182 | 1087 | 311 | 6958 | 3.1% |
| **5** | 301 | 1116 | 175 | 339 | 406 | 1167 | 1196 | 380 | 344 | 166 | 1015 | 255 | 6860 | 3.0% |
| **6** | 277 | 1141 | 158 | 343 | 435 | 1198 | 1206 | 406 | 327 | 169 | 1050 | 273 | 6983 | 3.1% |
| **7** | 256 | 1102 | 174 | 350 | 371 | 1174 | 1199 | 414 | 349 | 161 | 1143 | 273 | 6966 | 3.1% |
| **8** | 257 | 1122 | 169 | 337 | 403 | 1140 | 1222 | 365 | 341 | 164 | 1131 | 299 | 6950 | 3.1% |
| **9** | 282 | 1112 | 169 | 338 | 329 | 1147 | 1201 | 389 | 322 | 171 | 1053 | 261 | 6774 | 3.0% |
| **10** | 257 | 1182 | 148 | 360 | 369 | 1174 | 1137 | 415 | 323 | 159 | 1161 | 268 | 6953 | 3.1% |
| **11** | 269 | 996 | 167 | 305 | 377 | 1204 | 1159 | 354 | 311 | 170 | 1154 | 293 | 6759 | 3.0% |
| **12** | 247 | 1033 | 174 | 307 | 358 | 1174 | 1195 | 353 | 336 | 183 | 1127 | 275 | 6762 | 3.0% |
| **13** | 283 | 1045 | 185 | 321 | 364 | 1174 | 1178 | 388 | 293 | 200 | 1128 | 273 | 6832 | 3.0% |
| **14** | 267 | 1161 | 154 | 322 | 368 | 1164 | 1235 | 370 | 352 | 166 | 1065 | 258 | 6882 | 3.0% |
| **15** | 276 | 1117 | 167 | 321 | 391 | 1177 | 1174 | 361 | 297 | 168 | 1102 | 302 | 6853 | 3.0% |
| **16** | 290 | 1125 | 160 | 312 | 383 | 1175 | 1149 | 394 | 319 | 183 | 1095 | 269 | 6854 | 3.0% |
| **17** | 241 | 1115 | 188 | 331 | 387 | 1154 | 1249 | 372 | 318 | 183 | 1105 | 302 | 6945 | 3.1% |
| **18** | 279 | 1106 | 184 | 331 | 396 | 1185 | 1174 | 398 | 313 | 175 | 1126 | 241 | 6908 | 3.0% |
| **19** | 286 | 1074 | 167 | 318 | 420 | 1243 | 1141 | 364 | 322 | 165 | 1186 | 266 | 6952 | 3.1% |
| **20** | 255 | 1126 | 164 | 320 | 332 | 1131 | 1205 | 406 | 351 | 169 | 1127 | 275 | 6861 | 3.0% |
| **21** | 259 | 1114 | 150 | 326 | 398 | 1191 | 1133 | 446 | 333 | 181 | 1137 | 258 | 6926 | 3.1% |
| **22** | 281 | 1086 | 162 | 330 | 386 | 1088 | 1107 | 404 | 297 | 179 | 1136 | 282 | 6738 | 3.0% |
| **23** | 246 | 1129 | 173 | 327 | 382 | 1198 | 1235 | 359 | 335 | 183 | 1148 | 273 | 6988 | 3.1% |
| **24** | 285 | 1099 | 148 | 327 | 392 | 1201 | 1148 | 339 | 288 | 159 | 1071 | 269 | 6726 | 3.0% |
| **25** | 277 | 1113 | 165 | 312 | 392 | 1154 | 1157 | 377 | 299 | 167 | 1151 | 251 | 6815 | 3.0% |
| **26** | 272 | 1183 | 158 | 328 | 366 | 1137 | 1173 | 377 | 320 | 181 | 1092 | 286 | 6873 | 3.0% |

| | | | | | | | | | | | | | | |
|---|---|---|---|---|---|---|---|---|---|---|---|---|---|---|
| **27** | 273 | 1091 | 200 | 363 | 397 | 1130 | 1183 | 351 | 305 | 181 | 1223 | 246 | 6943 | 3.1% |
| **28** | 256 | 1111 | 171 | 343 | 411 | 1182 | 1139 | 408 | 328 | 159 | 1138 | 263 | 6909 | 3.0% |
| **29** | 278 | 1142 | 176 | 315 | 401 | 1135 | 1215 | 335 | 307 | 187 | 1074 | 291 | 6856 | 3.0% |
| **30** | 271 | 1139 | 193 | 309 | 376 | 1094 | 1154 | 395 | 318 | 176 | 1131 | 273 | 6829 | 3.0% |
| **31** | 282 | 1151 | 147 | 339 | 377 | 1110 | 1242 | 387 | 311 | 177 | 1140 | 269 | 6932 | 3.1% |
| **32** | 260 | 1143 | 155 | 314 | 439 | 1095 | 1177 | 355 | 351 | 183 | 1099 | 243 | 6814 | 3.0% |
| **33** | 260 | 1086 | 170 | 318 | 366 | 1192 | 1151 | 368 | 305 | 189 | 1084 | 263 | 6752 | 3.0% |
| **34** | 272 | 1146 | 158 | 310 | 368 | 1153 | 1203 | 369 | 314 | 178 | 1141 | 259 | 6871 | 3.0% |
| **35** | 284 | 1136 | 177 | 331 | 371 | 1092 | 1135 | 420 | 324 | 184 | 1105 | 282 | 6841 | 3.0% |
| **36** | 290 | 1121 | 176 | 332 | 329 | 1213 | 1144 | 363 | 322 | 171 | 1074 | 252 | 6787 | 3.0% |
| **37** | 267 | 1082 | 185 | 336 | 387 | 1214 | 1151 | 380 | 331 | 169 | 1147 | 272 | 6921 | 3.1% |
| **38** | 289 | 1162 | 167 | 307 | 375 | 1196 | 1199 | 383 | 309 | 151 | 1149 | 280 | 6967 | 3.1% |
| **39** | 271 | 1094 | 176 | 302 | 377 | 1147 | 1177 | 380 | 348 | 184 | 1085 | 254 | 6795 | 3.0% |
| **40** | 235 | 1097 | 151 | 333 | 381 | 1142 | 1139 | 406 | 328 | 161 | 1105 | 270 | 6748 | 3.0% |
| **41** | 282 | 1142 | 171 | 322 | 375 | 1093 | 1192 | 401 | 337 | 189 | 1119 | 280 | 6903 | 3.0% |
| **42** | 274 | 1155 | 155 | 350 | 358 | 1141 | 1148 | 368 | 313 | 162 | 1126 | 264 | 6814 | 3.0% |
| **43** | 256 | 1163 | 146 | 351 | 422 | 1262 | 1134 | 380 | 340 | 182 | 1097 | 292 | 7025 | 3.1% |
| **44** | 282 | 1136 | 168 | 358 | 382 | 1162 | 1195 | 370 | 323 | 159 | 1125 | 268 | 6928 | 3.1% |
| **45** | 256 | 1125 | 176 | 335 | 391 | 1246 | 1151 | 372 | 322 | 164 | 1076 | 280 | 6894 | 3.0% |
| **46** | 267 | 1065 | 173 | 335 | 382 | 1205 | 1199 | 390 | 330 | 188 | 1134 | 282 | 6950 | 3.1% |
| **47** | 305 | 1184 | 185 | 388 | 397 | 1131 | 1214 | 383 | 328 | 186 | 1112 | 258 | 7071 | 3.1% |
| **48** | 312 | 997 | 181 | 343 | 343 | 1064 | 1156 | 421 | 323 | 178 | 1157 | 270 | 6745 | 3.0% |
| **49** | 2146 | 587 | 102 | 198 | 178 | 625 | 612 | 2395 | 200 | 93 | 11006 | 130 | 18272 | 8.1% |
| **50** | 187 | 11457 | 116 | 196 | 2622 | 610 | 663 | 266 | 214 | 120 | 672 | 2278 | 19401 | 8.6% |
| **Total** | **4.4%** | **19.2%** | **2.1%** | **4.1%** | **5.9%** | **14.3%** | **14.5%** | **5.9%** | **4.0%** | **2.2%** | **19.1%** | **4.4%** | **391067** | **226655** |

**Supplementary Table 5:** Repartition of substitutions along the 50 bp probe sequence for mapped Infinium 450K probes on the *Macaca mulatta* genome. *Percentages are calculated over the total number of mapped probes (226,655), NOT the total number of substitutions (391,067) as one probe can have several mismatches.

| EPIC | *Macaca mulatta* | | | | | | | | | | | | | |
|---|---|---|---|---|---|---|---|---|---|---|---|---|---|---|
| **Base** | **A->C** | **A->G** | **A->T** | **C->A** | **C->G** | **C->T** | **G->A** | **G->C** | **G->T** | **T->A** | **T->C** | **T->G** | **ALL** | ***Percent** |
| **1** | 4887 | 1148 | 221 | 345 | 425 | 1128 | 1140 | 5542 | 374 | 266 | 27824 | 324 | 43624 | 10.8% |
| **2** | 284 | 25524 | 200 | 297 | 4427 | 1111 | 1091 | 300 | 335 | 202 | 1032 | 4270 | 39073 | 9.7% |
| **3** | 423 | 2195 | 331 | 531 | 675 | 2103 | 1873 | 556 | 570 | 284 | 1921 | 500 | 11962 | 3.0% |
| **4** | 460 | 2032 | 331 | 541 | 631 | 2041 | 2079 | 619 | 612 | 339 | 1918 | 498 | 12101 | 3.0% |
| **5** | 494 | 1970 | 356 | 584 | 625 | 2095 | 2151 | 637 | 597 | 330 | 1844 | 472 | 12155 | 3.0% |
| **6** | 453 | 2008 | 319 | 552 | 643 | 2130 | 2159 | 668 | 558 | 302 | 1966 | 459 | 12217 | 3.0% |
| **7** | 462 | 2001 | 335 | 574 | 611 | 2115 | 2115 | 648 | 606 | 313 | 2029 | 493 | 12302 | 3.1% |
| **8** | 439 | 1974 | 305 | 589 | 651 | 2073 | 2205 | 612 | 582 | 305 | 1995 | 472 | 12202 | 3.0% |
| **9** | 452 | 2001 | 320 | 574 | 560 | 2074 | 2148 | 618 | 579 | 310 | 1922 | 468 | 12026 | 3.0% |
| **10** | 435 | 2084 | 300 | 610 | 636 | 2153 | 2037 | 645 | 552 | 300 | 2081 | 458 | 12291 | 3.1% |
| **11** | 450 | 1890 | 331 | 564 | 645 | 2040 | 2108 | 604 | 534 | 327 | 1985 | 472 | 11950 | 3.0% |
| **12** | 453 | 1961 | 321 | 557 | 580 | 2088 | 2180 | 564 | 587 | 333 | 1989 | 485 | 12098 | 3.0% |
| **13** | 489 | 1882 | 348 | 567 | 597 | 2130 | 2060 | 638 | 514 | 350 | 1980 | 465 | 12020 | 3.0% |
| **14** | 445 | 1996 | 296 | 547 | 602 | 2119 | 2191 | 621 | 588 | 322 | 1933 | 442 | 12102 | 3.0% |
| **15** | 463 | 1972 | 316 | 589 | 628 | 2087 | 2163 | 596 | 527 | 311 | 1929 | 481 | 12062 | 3.0% |
| **16** | 493 | 1998 | 301 | 532 | 616 | 2060 | 2059 | 636 | 573 | 347 | 2001 | 459 | 12075 | 3.0% |
| **17** | 406 | 1972 | 350 | 564 | 623 | 2102 | 2170 | 599 | 566 | 346 | 2007 | 469 | 12174 | 3.0% |
| **18** | 458 | 2018 | 318 | 575 | 632 | 2163 | 2158 | 626 | 585 | 325 | 1993 | 419 | 12270 | 3.0% |
| **19** | 489 | 1901 | 328 | 541 | 637 | 2189 | 2057 | 584 | 531 | 325 | 2027 | 465 | 12074 | 3.0% |
| **20** | 460 | 2029 | 330 | 556 | 579 | 2084 | 2160 | 609 | 619 | 315 | 2028 | 456 | 12225 | 3.0% |
| **21** | 456 | 2021 | 296 | 549 | 647 | 2115 | 2144 | 665 | 543 | 325 | 1996 | 447 | 12204 | 3.0% |
| **22** | 463 | 2005 | 313 | 553 | 623 | 2019 | 1991 | 636 | 516 | 320 | 2019 | 484 | 11942 | 3.0% |
| **23** | 430 | 2019 | 334 | 558 | 619 | 2119 | 2206 | 578 | 558 | 314 | 2010 | 474 | 12219 | 3.0% |
| **24** | 467 | 1987 | 307 | 573 | 621 | 2200 | 2077 | 586 | 518 | 303 | 1977 | 462 | 12078 | 3.0% |
| **25** | 464 | 1989 | 312 | 544 | 657 | 2098 | 2073 | 629 | 554 | 318 | 2003 | 432 | 12073 | 3.0% |
| **26** | 461 | 2007 | 322 | 537 | 566 | 2033 | 2057 | 642 | 559 | 337 | 1944 | 463 | 11928 | 3.0% |

| | | | | | | | | | | | | | | |
|---|---|---|---|---|---|---|---|---|---|---|---|---|---|---|
| **27** | 456 | 1962 | 328 | 594 | 629 | 2020 | 2061 | 603 | 526 | 303 | 2169 | 414 | 12065 | 3.0% |
| **28** | 470 | 2009 | 328 | 581 | 656 | 2114 | 2098 | 644 | 545 | 313 | 2034 | 453 | 12245 | 3.0% |
| **29** | 474 | 2047 | 329 | 554 | 662 | 2064 | 2125 | 568 | 548 | 349 | 1917 | 488 | 12125 | 3.0% |
| **30** | 469 | 2046 | 333 | 562 | 611 | 2073 | 2121 | 628 | 566 | 311 | 1999 | 491 | 12210 | 3.0% |
| **31** | 472 | 2027 | 300 | 579 | 639 | 2058 | 2174 | 609 | 560 | 354 | 2033 | 479 | 12284 | 3.0% |
| **32** | 426 | 2049 | 285 | 560 | 673 | 2038 | 2120 | 603 | 579 | 331 | 1983 | 446 | 12093 | 3.0% |
| **33** | 461 | 1983 | 332 | 557 | 585 | 2168 | 2110 | 605 | 543 | 351 | 1994 | 438 | 12127 | 3.0% |
| **34** | 445 | 1988 | 309 | 565 | 608 | 2139 | 2151 | 585 | 566 | 308 | 2055 | 463 | 12182 | 3.0% |
| **35** | 491 | 2038 | 312 | 578 | 589 | 2037 | 2083 | 637 | 570 | 329 | 1966 | 488 | 12118 | 3.0% |
| **36** | 466 | 2018 | 322 | 568 | 580 | 2096 | 2045 | 609 | 560 | 328 | 1973 | 455 | 12020 | 3.0% |
| **37** | 454 | 1878 | 321 | 561 | 648 | 2186 | 2060 | 609 | 566 | 319 | 2026 | 464 | 12092 | 3.0% |
| **38** | 452 | 2068 | 313 | 544 | 644 | 2129 | 2156 | 619 | 536 | 296 | 2020 | 463 | 12240 | 3.0% |
| **39** | 456 | 1970 | 324 | 551 | 610 | 2087 | 2118 | 609 | 599 | 329 | 1982 | 451 | 12086 | 3.0% |
| **40** | 455 | 2011 | 313 | 560 | 624 | 2066 | 2060 | 653 | 606 | 305 | 1996 | 445 | 12094 | 3.0% |
| **41** | 452 | 2115 | 343 | 601 | 607 | 1997 | 2145 | 609 | 586 | 323 | 1967 | 474 | 12219 | 3.0% |
| **42** | 454 | 2054 | 322 | 586 | 596 | 2123 | 2055 | 617 | 571 | 308 | 2006 | 458 | 12150 | 3.0% |
| **43** | 450 | 1980 | 287 | 608 | 664 | 2170 | 2080 | 605 | 587 | 346 | 2027 | 485 | 12289 | 3.0% |
| **44** | 451 | 2011 | 321 | 610 | 639 | 2065 | 2171 | 598 | 537 | 320 | 1985 | 473 | 12181 | 3.0% |
| **45** | 430 | 1936 | 321 | 615 | 666 | 2202 | 2022 | 621 | 524 | 297 | 1971 | 479 | 12084 | 3.0% |
| **46** | 478 | 1935 | 309 | 583 | 589 | 2148 | 2115 | 619 | 561 | 360 | 2079 | 456 | 12232 | 3.0% |
| **47** | 506 | 2041 | 324 | 650 | 641 | 2063 | 2136 | 606 | 559 | 362 | 2037 | 452 | 12377 | 3.1% |
| **48** | 532 | 1915 | 338 | 569 | 577 | 1973 | 2125 | 692 | 540 | 368 | 2140 | 438 | 12207 | 3.0% |
| **49** | 4422 | 1051 | 176 | 347 | 317 | 1084 | 1174 | 4452 | 334 | 208 | 25474 | 257 | 39296 | 9.8% |
| **50** | 329 | 28008 | 228 | 329 | 5565 | 1118 | 1189 | 417 | 369 | 225 | 1145 | 4935 | 43857 | 10.9% |
| **Total** | 4.3% | 20.4% | 2.1% | 3.8% | 5.4% | 13.9% | 14.0% | 5.4% | 3.8% | 2.2% | 20.3% | 4.3% | 724320 | 402935 |

**Supplementary Table 7:** Repartition of substitutions along the 50 bp probe sequence for mapped Infinium EPIC probes on the *Macaca mulatta* genome. *Percentages are calculated over the total number of mapped probes (402,935), NOT the total number of substitutions (724,320) as one probe can have several mismatches.

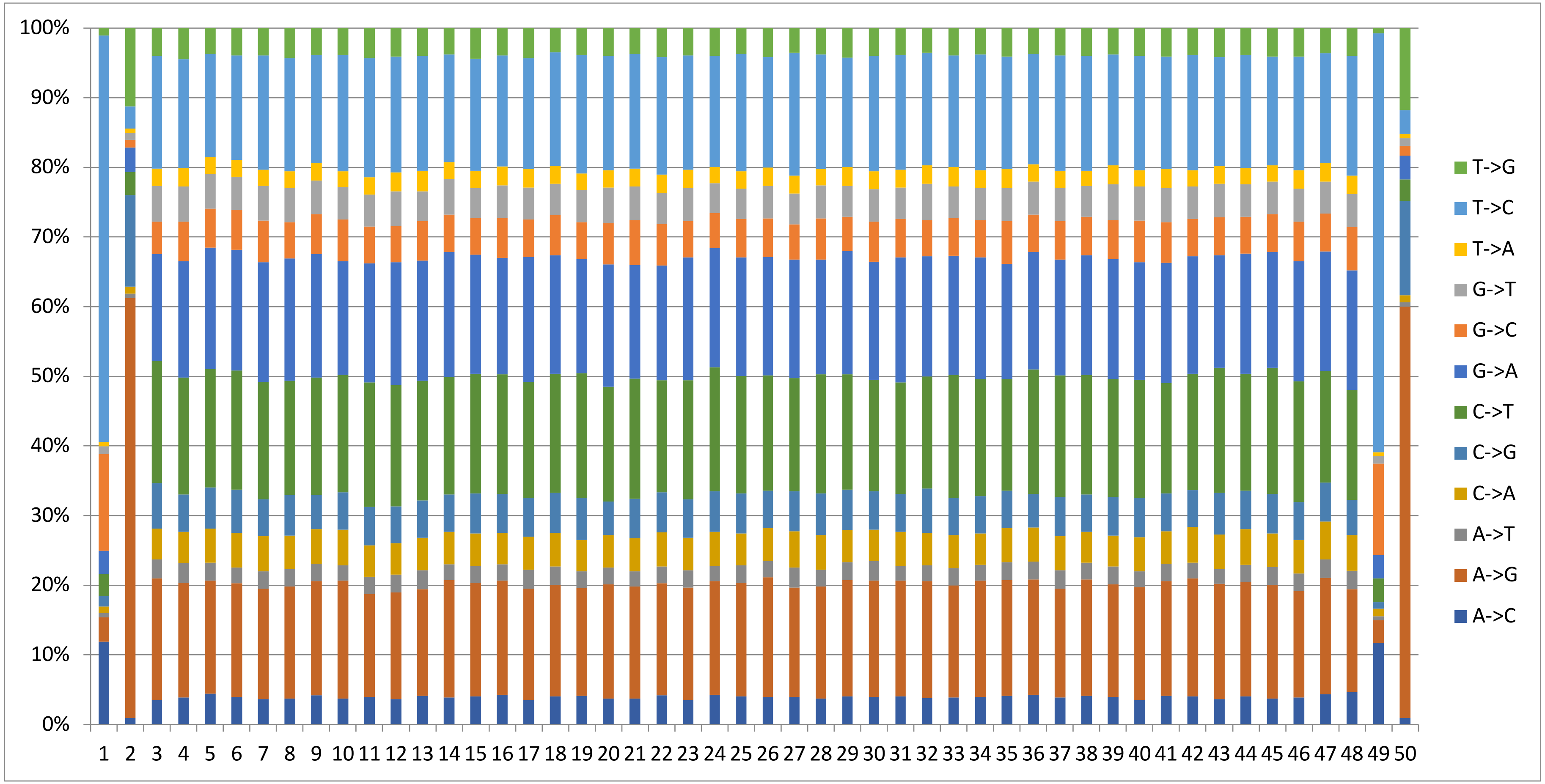


**Suppl. Figure 1:** Repartition of substitutions along the 50 bp of the human-designed probe sequences of the Infinium 450K mapped to the *Macaca mulatta* genome.
